# *Vibrio cholerae* colonization of the arthropod intestine activates innate immune signaling in enteroendocrine cells via phosphorylation of the nuclear receptor ultraspiracle

**DOI:** 10.1101/2025.04.21.649885

**Authors:** Xiang Ding, Paula I. Watnick

## Abstract

The Gram-negative rod *Vibrio cholerae* causes profuse diarrhea in humans and is found in close association with both terrestrial and aquatic arthropods in the environment. We have previously shown that *V. cholerae* colonizes the arthropod intestine. Here we show that tryptophan produced by *V. cholerae* in the arthropod intestine is used by enterocytes to synthesize serotonin and signal to enteroendocrine cells (EECs) to activate the TNF-like immune deficiency pathway IMD. We define the EEC serotonin signaling pathway, which involves a subset of serotonin G protein-coupled receptors, Gαq, phospholipase C, and protein kinase C. This pathway culminates in phosphorylation and potentiation of the nuclear receptor ultraspiracle, which binds the sex hormone ecdysone to activate IMD signaling. While IMD signaling increases antimicrobial peptide synthesis in all intestinal cell types, IMD signaling in EECs uniquely activates expression of the enteroendocrine peptide Tachykinin and the *V. cholerae* colonization factor Peritrophin-15a. We propose that, because *V. cholerae* secretes a metabolite that activates IMD signaling in EECs, it has evolved to exploit the arthropod intestinal innate immune response to maximize adhesion to the arthropod intestine.

## Introduction

The epithelia of animals are bombarded with bacterial fragments, colonized by non-invasive commensal microbes, and challenged by pathogens. Discernment of the context of these microbial encounters is aided by epithelial G-protein coupled receptors (GPCRs) that respond to microbial metabolites and pattern recognition receptors that bind microbe-associated molecular patterns (MAMPs) such as peptidoglycan (PG) or lipopolysaccharide. These tissues must then deploy an appropriate innate immune response to each microbial encounter ^1, 2, 3, 4^.

The ability to detect microbes and respond appropriately is particularly critical to the intestinal innate immune response. The constant flow of nutrients in the intestine promotes rapid microbial growth leading to persistent colonization by commensal organisms and sporadic colonization by pathogens. In this nutrient-rich environment, microbial metabolites such as short chain fatty acids and amino acids play a particularly important role in modulating intestinal physiology, metabolism, and immune activation ^5, 6, 7^.

*Vibrio cholerae* is a Gram-negative bacterium, a human diarrheal pathogen, and a natural inhabitant of aquatic environments ^8, 9^. Many studies suggest that *V. cholerae* is associated with terrestrial insects and zooplankton in these environments, and the chitinous surfaces of these organisms provide both nutrition and a signal that activates *V. cholerae* natural competence ^10, 11, 12, 13, 14, 15, 16^. While we use the *Drosophila* intestine as a model for the interaction of the human pathogen *V. cholerae* with the intestines of terrestrial and aquatic arthropods, this model has also informed the behavior of *V. cholerae* in intestinal environment, its interaction with commensal microbes, and the response of intestinal cells ^17, 18, 19, 20, 21^.

The intestine of an adult fly has three compartments: the foregut, midgut and hindgut. The midgut, which is the subject of this study, functions in nutrient absorption, commensal control, and pathogen defense ^22, 23^. It is further subdivided into the anterior (AMG), middle (MMG) and posterior (PMG) midgut regions, each of which has a distinct function and morphology. The midgut epithelium is lined by a mucus-like membrane termed the peritrophic matrix (PM). The PM consists of a network of chitin fibrils with associated chitin-binding proteins that include the peritrophins ^24, 25^. Within the AMG, three morphological regions R0, R1, and R2 can be distinguished ^23^. The distal part of the AMG, which abuts the MMG, is designated R2.

The *Drosophila* AMG is comprised of three mature cell types, namely enterocytes (ECs), enteroendocrine cells (EECs), and intestinal stem cells (ISCs). Approximately 90% of stem cells will ultimately differentiate to enterocytes, which make up most of the epithelium ^26^. In addition to their function as absorptive cells, enterocytes sense and respond to microbial metabolites such as uracil and microbial components such as PG by producing two types of bactericidal molecules, reactive oxygen species (ROS) and anti-microbial peptides (AMPs) ^27, 28, 29^. Enteroendocrine cells, whose differentiation from ISCs is dependent on the transcription factor *prospero* (pros), are a rare cell type in intestinal epithelium. EEC populations can be divided into two major groups based on the peptide (EEPs) hormones they express. Class I EECs express Allatostatin C (AstC), while class II EECs express Tachykinin (Tk) ^30^. These two classes have unique functions in host physiology.

Every cell type in the *Drosophila* intestine expresses components of the immune deficiency (IMD) pathway, which is homologous to the TNF pathway of mammals ^31^. IMD signaling is triggered by meso-diaminopimelic acid (DAP)-type peptidoglycan (PGN), which comprises the cell wall of Gram-negative and some Gram-positive bacteria. PGRP-LC and PGRP-LE are the IMD-specific pattern recognition receptors (PRRs) that bind PGN. PGRP-LC is a trans-membrane receptor that binds extracellular PG polymers, while PGRP-LE, an intracellular receptor, specifically binds monomeric PG ^32^. The activation of these receptors results in cleavage and phosphorylation of the transcription factor Relish (Rel) by recruiting two complexes, complex I consisting of IMD, the adaptor protein dFadd and the caspase-8 homolog Dredd and complex II consisting of I-κB kinase ϒ homolog Kenny (Key) and I-κB kinase β. The active N-terminus of Rel translocates to nucleus where it promotes transcription of genes coding for AMPs, such as Diptericin A (DptA), Attacin A (AttA), and Cecropin A1 (CecA1) as well as cell-type specific transcripts.

Our recent research has shown that the IMD pathway in Tk+ EECs of the AMG is activated by metabolites such as the short-chain fatty acid acetate ^33, 34^. In these cells, IMD signaling promotes transcription of *Tk* in addition to IMD-specific AMPs. Because ECs are a major source of some AMPs, this suggests that Tk promotes innate immune crosstalk between epithelial cells ^31^. However, the mechanism by which this response is coordinated remains unexplored.

In mammals, the amino acid tryptophan is a hub for communication between microbes and the host intestine ^35^. It is the least abundant amino acid in both microbial and animal proteins, accounting for approximately 1% of amino acids incorporated into cytoplasmic proteins ^36^. As a result, tryptophan is ideal for use in non-structural applications such as energy generation, host-microbe communication, and synthesis of signaling molecules. While many bacteria possess the enzymes necessary for its synthesis, tryptophan is an essential amino acid for all animals.

Microbial synthesis of tryptophan is costly and, therefore, tightly controlled at the transcriptional level^37^. When tryptophan is abundant in the environment, microbes prioritize tryptophan import ^38^. Tryptophan not needed for protein synthesis is then catabolized to indole and pyruvate ^38, 39, 40^. While pyruvate enters the tricarboxylic acid cycle, indole is excreted. In the mammalian intestine, indole is sensed by the arylhydrocarbon receptor ^41, 42^. Because the diet is believed to be a major source of tryptophan in the mammalian intestine, little attention has been paid to the fate of microbially synthesized tryptophan that is secreted into the gut ^43^.

We previously showed that avid bacterial consumption of dietary tryptophan in the arthropod intestine quenches the intestinal immune response ^44^. Here we show that wild-type *V. cholerae* actively promotes the innate immune response of the arthropod intestine by providing ECs with tryptophan for serotonin synthesis. Serotonin activates IMD pathway signaling in Tk+ EECs. This increases expression of the small host PM protein Peritrophin 15a, which is a colonization factor for *V. cholerae* ^45^. We define the complete signaling pathway leading from activation of serotonin receptors in Tk+ EECs to activation of protein kinase C, to phosphorylation of the nuclear ecdysone sex hormone receptor ultraspiracle (usp). This increases expression of the PRR PGRP-LC in Tk+ EECs, which enhances IMD signaling. We propose that *V. cholerae*, which likely has evolved with arthropods, recruits EECs to the host intestinal innate immune response to enhance its own colonization.

## Results

### *V. cholerae* tryptophan synthesis activates serotonin production in ECs to increase numbers of Tk+ EECs and AMP transcription

In the first and rate-limiting step of serotonin synthesis, tryptophan is converted to 5-hydroxytryptophan (5-HTP) by tryptophan hydroxylase, which is encoded by the *Drosophila* gene *Henna* (Fig 1A). 5-HTP is converted to serotonin (5-hydroxytryptamine, 5-HT) by 5-HTP decarboxylase (Ddc). By studying *Drosophila* infection with the *V. cholerae* mutant Δ*hapR*, we previously discovered that bacterial consumption of tryptophan abolishes serotonin synthesis in the *Drosophila* intestine ^44^. Because diet is reported to be a major source of intestinal tryptophan, we did not investigate the impact of WT *V. cholerae* on intestinal serotonin synthesis ^46^. To explore this, we first questioned whether colonization with WT *V. cholerae* modulates *Hn* expression. We observed a significant increase in Henna expression in the intestines of Oregon R (OreR) flies infected with wild-type (WT) *V. cholerae* as compared with uninfected flies (Fig 1B). Immunofluorescence using an anti-5-HT antibody showed increased serotonin levels in the intestines of *V. cholerae-*infected flies (Fig 1C and D). While most of the 5-HT was located outside of EECs that express Tk (Tk+ EECs), some also co-localized with Tk+ EECs (Fig 1C). Because serotonin can be imported by the transporter SerT and no serotonin co-localized with Tk+ cells in the absence of *V. cholerae* infection, it is most likely that some of the serotonin synthesized by ECs is imported into EECs and packaged ^31, 47^. We conclude that WT *V. cholerae* infection promotes intestinal serotonin synthesis by inducing *Henna* expression.

**Figure 1:**
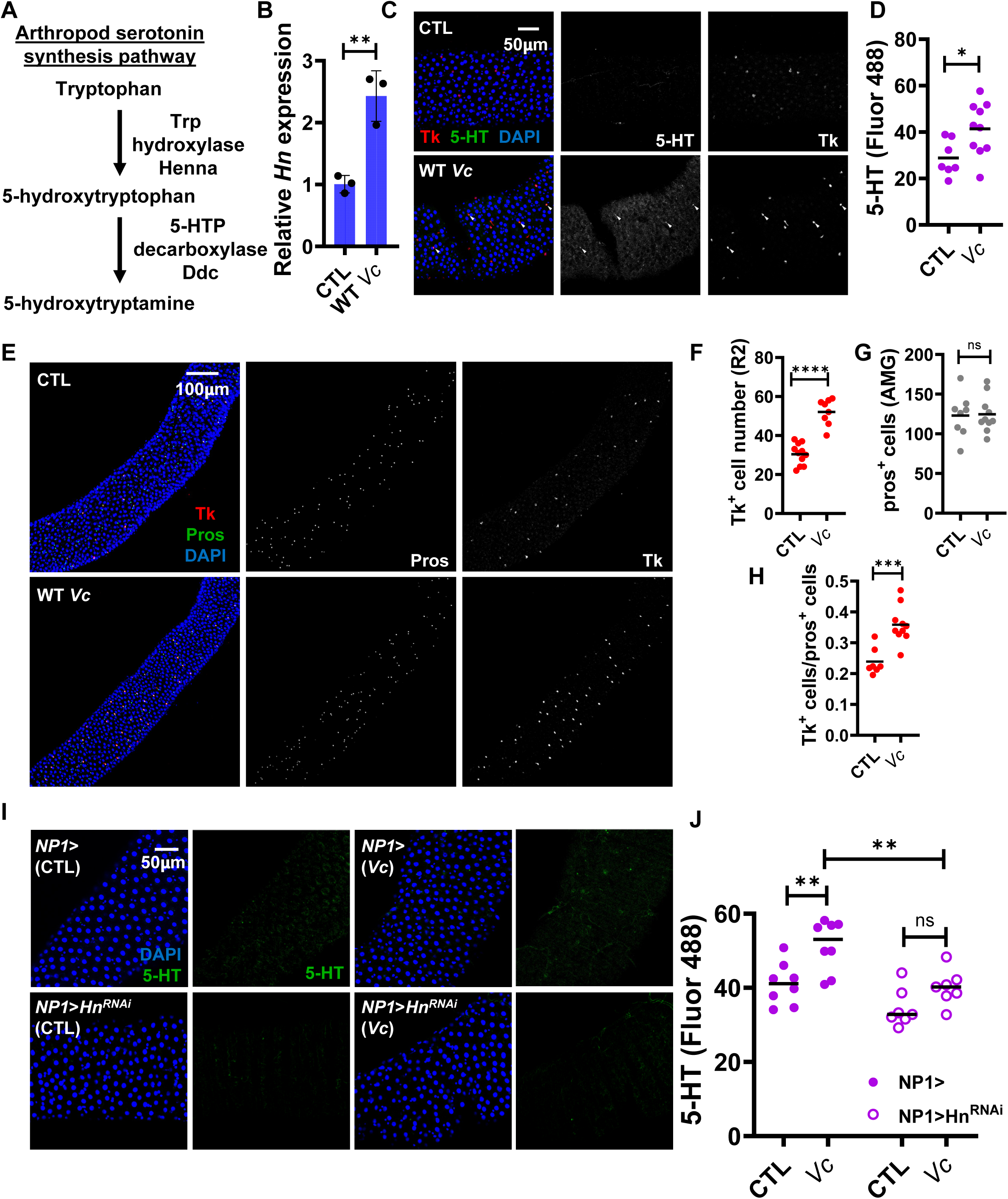
*V. cholerae* infection activates serotonin (5-HT) synthesis in the arthropod intestine leading to an increase in Tachykinin (Tk)+ cells, see also Figure S1. (A) The arthropod serotonin synthesis pathway from tryptophan, which is an essential amino acid. (B) RT-qPCR of *Henna* (*Hn*), which encodes a tryptophan hydroxylase in uninfected (CTL) and *V. cholerae*-infected (*Vc*) Oregon R flies. Bars represent the mean of biological triplicates. A student’s t test was used to assess significance. (C) Micrographs and (D) quantification of 5-HT immunofluorescence of the AMG of control flies and flies infected with *Vc*. Tk immunofluorescence is also shown. Arrows indicate 5-HT co-localization with Tk+ EECs. The mean of six intestinal measurements is shown. A student’s t test was applied to assess significance. (E) Micrographs and (F-H) quantification of Tk+ and Prospero (Pros) + cells in the AMG. A student’s t test was used to assess significance. (I) Micrographs and (J) quantification of 5-HT immunofluorescence in the intestines of NP1> enterocyte driver-only flies and NP1>*Hn*^RNAi^ flies. Flies were either uninfected (CTL) or infected with *V. cholerae* (Vc). The mean of at least six intestines is shown. An ordinary one-way ANOVA with Tukey’s multiple comparisons test was used to assess significance. Error bars represent the standard deviation. **** p<0.0001, *** p<0.001, ** p<0.01, * p<0.05, ns not significant.

Tryptophan consumption by a *V. cholerae* Δ*hapR* mutant decreases the numbers of Tk+ EECs ^44^. We questioned whether infection with WT *V. cholerae* would increase Tk+ cells. Compared with uninfected flies, *V. cholerae* infection significantly increased the Tk+ cell number in the R2 region of the AMG but not in the PMG (Fig 1E-F and Fig S1). To rule out the possibility that this reflected a general increase in EECs, we marked all EECs with an anti-Prospero antibody. The total number of EECs remained constant with *V. cholerae* infection, while the proportion of EECs that expressed Tk increased (Fig 1G and H). This indicates that *V. cholerae* infection increases Tk+ EECs in the arthropod midgut.

To identify the cell type responding to *V. cholerae* infection, we blocked serotonin synthesis in ECs by expressing *Hn*^RNAi^ driven by NP1-Gal4 (NP1>). In NP1> driver control flies, infection significantly increased serotonin synthesis. This was reduced by NP1>*Hn*^RNAi^ in infected flies, and a comparison between NP1>*Hn*^RNAi^ -infected and uninfected flies did not reveal a significant difference in serotonin synthesis (Fig 1I and J). These data suggest that *V. cholerae* infection stimulates serotonin synthesis through activation of *Hn* transcription in enterocytes.

### Evidence that *V. cholerae* secretes tryptophan to modulate the intestinal innate immune response

*V. cholerae* synthesizes tryptophan from chorismate through the pathway shown in Fig 2A. To determine whether *V.* cholerae-synthesized tryptophan is responsible for activation of host serotonin synthesis in response to *V. cholerae* infection, we compared intestinal serotonin synthesis in WT *V cholerae* and a *V. cholerae* Δ*trpE* mutant in which the tryptophan synthesis pathway is blocked. Infection with the *V. cholerae* Δ*trpE* mutant did not increase serotonin synthesis nor Tk+ EECs in the intestine (Fig 2B-D). In the absence of provision of tryptophan by *V. cholerae,* intestinal *Hn* transcription was not significantly different from that of uninfected flies (Fig 2E). While transcription of IMD-activated AMPs was decreased in flies infected with the Δ*trpE* mutant, they remained significantly higher than that of uninfected flies (Fig 2F). This suggests that other bacterial products besides tryptophan, perhaps such as PGN, also contribute to the immune response observed during *V. cholerae* infection. Bacterial tryptophan synthesis and uptake are tightly controlled at the transcriptional level ^44^. We considered the possibility that a Δ*trpE* mutant might decrease intestinal serotonin synthesis and innate immune signaling by causing upregulation of the *V. cholerae* tryptophan transporters and tryptophan uptake. In other words, our data could also be consistent with removal of tryptophan from the intestine by a Δ*trpE* mutant rather than provision of tryptophan by WT *V. cholerae.* To examine this possibility, we measured transcription of the *V. cholerae* transporters *trpT* and *aaxT* in the intestines of flies infected with either WT *V. cholerae* or a Δ*trpE* mutant. To assess the accuracy of our measurement, we also measured transcription of *trpE*. As shown in Figure 2G, no significant difference was observed in transcription of the transporters, while *trpE* was significantly decreased in a Δ*trpE* mutant. We conclude that *V. cholerae* provides tryptophan to the host intestine to augment serotonin synthesis and activate the innate immune response.

**Figure 2:**
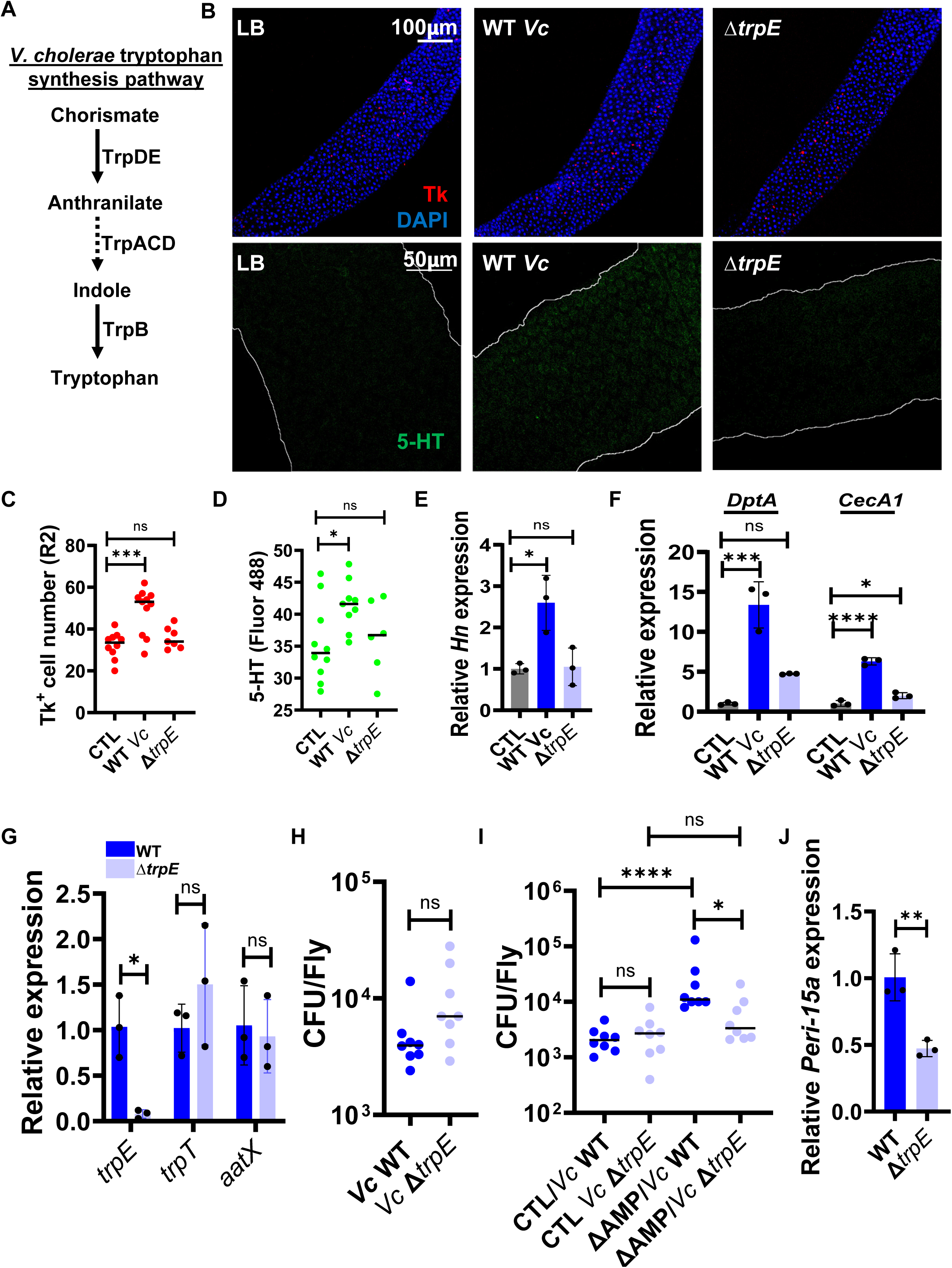
*V. cholerae-*synthesized tryptophan is converted to serotonin by enterocytes, which activates the innate immune response. (A) Schematic showing the *V. cholerae* tryptophan synthesis pathway. (B) Immunofluorescent images and quantification of (C) Tk+ cells and (D) 5-HT staining in Oregon R control flies fed LB alone or inoculated with wild-type *V. cholerae* (WT *Vc*) or a Δ*trpE V. cholerae* mutant. The brightness of the 5-HT micrographs was similarly corrected to improve visualization. The mean of at least six intestines is shown for each condition. An ordinary one-way ANOVA was used to assess significance. RT-qPCR analysis of (E) *Hn* and (F) *DptA* and *CecA1* expression. Bars represent the mean of three biological replicates. An ordinary one-way ANOVA was used to assess significance. (G) RT-qPCR analysis of *V. cholerae* tryptophan synthesis gene *trpE* and the *V. cholerae* tryptophan import genes *trpT* and *aaxT* in the intestines of OreR flies infected with either WT *V. cholerae* or a Δ*trpE* mutant. Bars represent the mean of three biological replicates. A student’s t test was used to assess significance. (H) Burden of WT *V.cholerae* and a Δ*trpE* mutant in the intestines of infected OreR flies. The mean burden of 8 flies is shown. Data were log-transformed and a Mann-Whitney test was used to assess significance of the transformed data. (I) Burden of WT *V.cholerae* and a Δ*trpE* mutant in the intestines of infected parental (CTL) flies or flies with 10 AMPs deleted (ΔAMP). Data were log-transformed, and a one way ANOVA with Tukey’s multiple comparisons test was used to assess significance. (J) RT-qPCR analysis of *Peri-15a* in the intestines of OreR flies infected with either WT *V. cholerae* or a Δ*trpE* mutant. Bars represent the mean of three biological replicates. A student’s t test was used to assess significance. Where shown, error bars represent the standard deviation. *** p<0.001, ** p<0.01, * p<0.05, ns not significant.

### *V. cholerae* tryptophan synthesis enhances intestinal colonization by activating expression of the colonization factor Peritrophin-15a (Peri-15a)

Synthesis and secretion of tryptophan is energetically costly. We questioned, therefore, whether this might provide *V. cholerae* with an advantage in colonization of the arthropod intestine. Interestingly, despite the increase in transcription of AMPs, colonization of the intestines of Ore R flies by WT *V. cholerae* and a Δ*trpE* mutant were not significantly different (Fig 2H). We reasoned either (i) that *V. cholerae* is resistant to the action of AMPS or (ii) that, in addition to the canonical IMD AMP response, *V. cholerae* tryptophan synthesis activates expression of a host colonization factor. To test this, we compared WT *V. cholerae* and Δ*trpE* mutant colonization of a parental *w*^1118^ strain with one carrying a deletion of 10 AMPs (ΔAMP) ^48^. Again, colonization of parental flies by a Δ*trpE* mutant was not significantly different from that by WT *V. cholerae* (Fig 2I). However, WT *V. cholerae* colonized the intestines of ΔAMP flies significantly better than those of the parental line, suggesting that *V. cholerae* is, in fact, susceptible to the AMPs produced in the fly intestine. Furthermore, in the absence of a contribution from AMPS, WT *V. cholerae* colonized the fly intestine significantly better than the Δ*trpE* mutant. Because tryptophan synthesis is not essential for colonization of the host intestine, we hypothesized that *V. cholerae* tryptophan synthesis might activate a host colonization factor. We previously identified Peritrophin-15a (Peri-15a), a component of the host peritrophic matrix, as a host colonization factor regulated by Tk ^45^. To determine if *Peri-15a* was differentially regulated in response to *V. cholerae*-produced tryptophan, we measured *Peri-15a* transcription in the intestines of flies infected with either WT *V. cholerae* or a Δ*trpE* mutant. As shown in Fig 2J, transcription of *Peri-15a* was decreased in the Δ*trpE* mutant. This suggests that tryptophan secretion by *V. cholerae* may represent an adaptation that increases colonization of the arthropod host intestine by promoting expression of Peri-15a.

### In uninfected flies, serotonin synthesized in ECs plays an important role in IMD signaling, Tk expression, and maintenance of lipid homeostasis

All animals are tryptophan auxotrophs. Therefore, uninfected flies must also have a source of tryptophan. In laboratory-raised flies, tryptophan is supplied by a nutrient-optimized diet. When tryptophan in the diet is limited, the slowed development of germ-free larvae can be rescued by the commensal *Acetobacter plantarum* ^49^. This suggests that *A. plantarum* may also provide tryptophan to the fly. Although the level of serotonin synthesis in the intestines of uninfected flies was below the level of detection by immunofluorescent staining, the literature suggested that dietary or microbiota-provided tryptophan might be available to serve as a substrate for basal serotonin synthesis. Because we could not detect serotonin in the intestines of uninfected flies by immunofluorescence, we instead investigated the phenotype of *Hn*^RNAi^ flies. Under homeostatic conditions, the majority of intestinal serotonin is synthesized in EECs known as enterochromaffin cells in mammals ^50^. We, therefore, measured transcription of AMPs in the intestines of uninfected Tk> and *Tk*>*Hn^RNAi^* flies. AMP transcription was indistinguishable in these flies suggesting that serotonin synthesis in Tk+ EECs is negligible (Fig S2A). In contrast*, Hn*^RNAi^ in ECs significantly decreased AMP expression (Fig S2B). We also observed reduced Tk+ EECs in the AMG of NP1*>Hn^RNAi^* flies and increased lipid accumulation, which results from insufficient Tk expression (Fig S2C and D) ^51^. We previously showed that acetate supplied by the microbiota is essential for maintenance of innate immune signaling, Tk expression and appropriate intestinal lipid metabolism under homeostatic conditions ^33, 34^. The results we present here demonstrate that synthesis of serotonin and serotonin signaling is also essential for maintenance of intestinal lipid homeostasis.

### Supplementation of the diet of uninfected flies with the serotonin precursor 5-HTP further increases Tk and AMP expression

To assess the impact of augmentation of serotonin synthesis on the innate immune response independent of infection and, therefore, MAMPs, we fed files Luria-Bertani broth (LB) alone or supplemented with the serotonin precursor 5-HTP and determined whether this was sufficient to increase serotonin levels in the gut. As shown in Fig S3A and B, 5-HTP supplementation increased intestinal serotonin levels significantly. We then asked whether 5-HTP supplementation affected Tk and AMP expression. This treatment increased both numbers of Tk+ cells in the R2 region of the AMG and expression of the IMD-controlled AMPs CecA1 and AttA (Fig S3C-E). As we observed during *V. cholerae* infection, the increase in Tk+ cells was not the result of an overall increase in EECs (Fig S3F and G). We reasoned that dietary supplementation with 5-HTP might affect the innate immune response by altering the microbiota. The microbiota of *Drosophila* raised in the laboratory consists almost exclusively of *Acetobacter* and *Lactobacillus* species ^52^. To show that 5-HTP-induced IMD activation was not driven by an increase in the microbiota, we measured the number of *Acetobacter* and *Lactobacillus* sp in the intestines of *Drosophila* fed LB alone or supplemented with 5-HTP. The microbial load and composition remained unchanged (Fig S3H). These results suggest that 5-HTP supplementation is sufficient to increase serotonin synthesis, Tk expression, and the innate immune response independent of a change in the microbiota or *V. cholerae* infection. As a final test of the validity of using 5-HTP supplementation to study serotonin signaling in the gut, we confirmed that tryptophan supplementation could also increase Tk+ EECs (Fig S3 I and J). Furthermore, in NP1>*Hn*^RNAi^ flies, numbers of Tk+ EECs were rescued by 5-HTP, which bypasses the requirement for Hn but not by tryptophan, which is a substrate of Hn. Based on these experiments, we conclude that serotonin synthesis in ECs is both necessary and sufficient to activate Tk expression and the intestinal innate immune response independent of infection. This provides us with a diet-based model to explore the serotonin signaling pathway in EECs independent of infection.

### Serotonin activates G-protein coupled receptor (GPCR) signaling in Tk+ cells

Serotonin binds GPCRs found on the surface of many intestinal cells. The *Drosophila* genome encodes 5 serotonin receptors: 5-HT1A,5-HT1B, 5-HT2A, 5-HT2B, 5-HT7 ^53^. To identify the GPCRs responding to serotonin on Tk+ cells, we knocked these down singly using gene-specific RNAi and the Tk-Gal4 driver and found that three GPCRs, 5-HT1A,5-HT1B, 5-HT7, decreased Tk+ EECs and increased lipid accumulation in the AMG of flies (Fig 3A and B).

**Figure 3:**
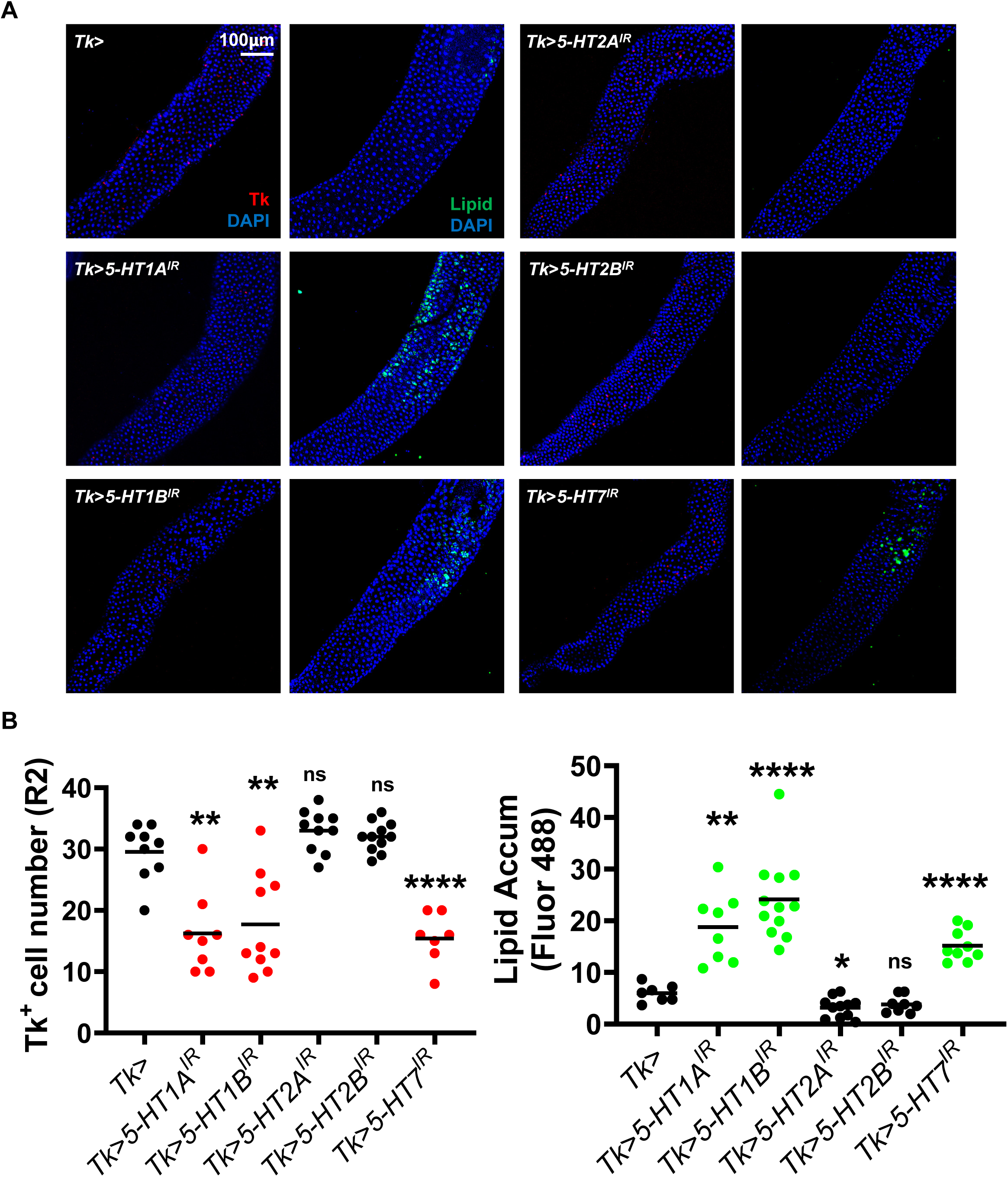
The receptors 5-HT1A,1B, and 7 participate in the enteroendocrine response to serotonin. (A) Immunofluorescent images and (B) quantification of Tk+ cells and Lipid (Bodipy) staining in the AMG (R2) of control Tk> driver-only flies or the indicated Tk>5-HT^RNAi^ receptor knockdown flies. The mean of the intestines of at least six flies is shown. Welch’s and Brown-Forsythe ANOVAs were used to assess significance. **** p<0.0001, ** p<0.01, ns not significant

As GPCRs, 5-HT receptors activate intracellular signaling cascades through heterotrimeric G-proteins with subunits Gα, Gβ, Gϒ ^54^. Gα proteins are divided into the four subtypes Gαi, Gαs, Gαq, and Gαo, each of which targets a specific signaling cascade ^54, 55^. Gαi and Gαo decrease intracellular cyclic adenosine monophosphate (cAMP) levels by inhibiting the cAMP synthase adenylate cyclase (AC), whereas Gαs increases cAMP levels by activating AC ^56^. Gαq mediates phospholipase C (PLC) activation. To define the signaling pathways downstream of the 5-HT receptors in Tk+ EECs, we knocked down all Gα subtypes using the Tk-Gal4 driver. The AMG of Tk>*Gαo*^RNAi^ and Tk> *Gαq*^RNAi^ flies showed decreased Tk+ EECs and increased lipid accumulation (Fig 4A and B and Fig S4A and B). GPCRs other than serotonin receptors could also regulate Tk+ expression via their interaction with Gα subunits. To determine whether Gαo and Gαq, in fact, function downstream of serotonin, we tested whether Tk>*Gαo*^RNAi^ and Tk> *Gαq*^RNAi^ as well as other Gα subtypes could block an increase in Tk+ EECs in response to 5-HTP. Only depletion of Gαq in Tk+ EECs blocked an increase in Tk expression (Fig 4A and B). This suggests that Gαq functions downstream of 5-HT receptors, while Gαo functions downstream of a different GPCR to increase Tk expression. As Gαq regulates PLC activation, we next queried the function of PLC in Tk+ EECs. There are 3 PLC paralogs encoded in fly genome, namely phospholipase 21C (Plc21C), no receptor potential (norpA), and small wing (sl). Only Tk>*norpA*^RNAi^ and Tk>*sl*^RNAi^ increased lipid accumulation in the gut and prevented 5-HTP-induced Tk expression, suggesting that these PLCs have overlapping functions in serotonin signaling (Fig 4C and D and S4C and D). These results show that the Gαq-PLC pathway acts downstream of serotonin and the 5-HT receptors to regulate Tk expression and lipid homeostasis in the *Drosophila* intestine.

**Figure 4:**
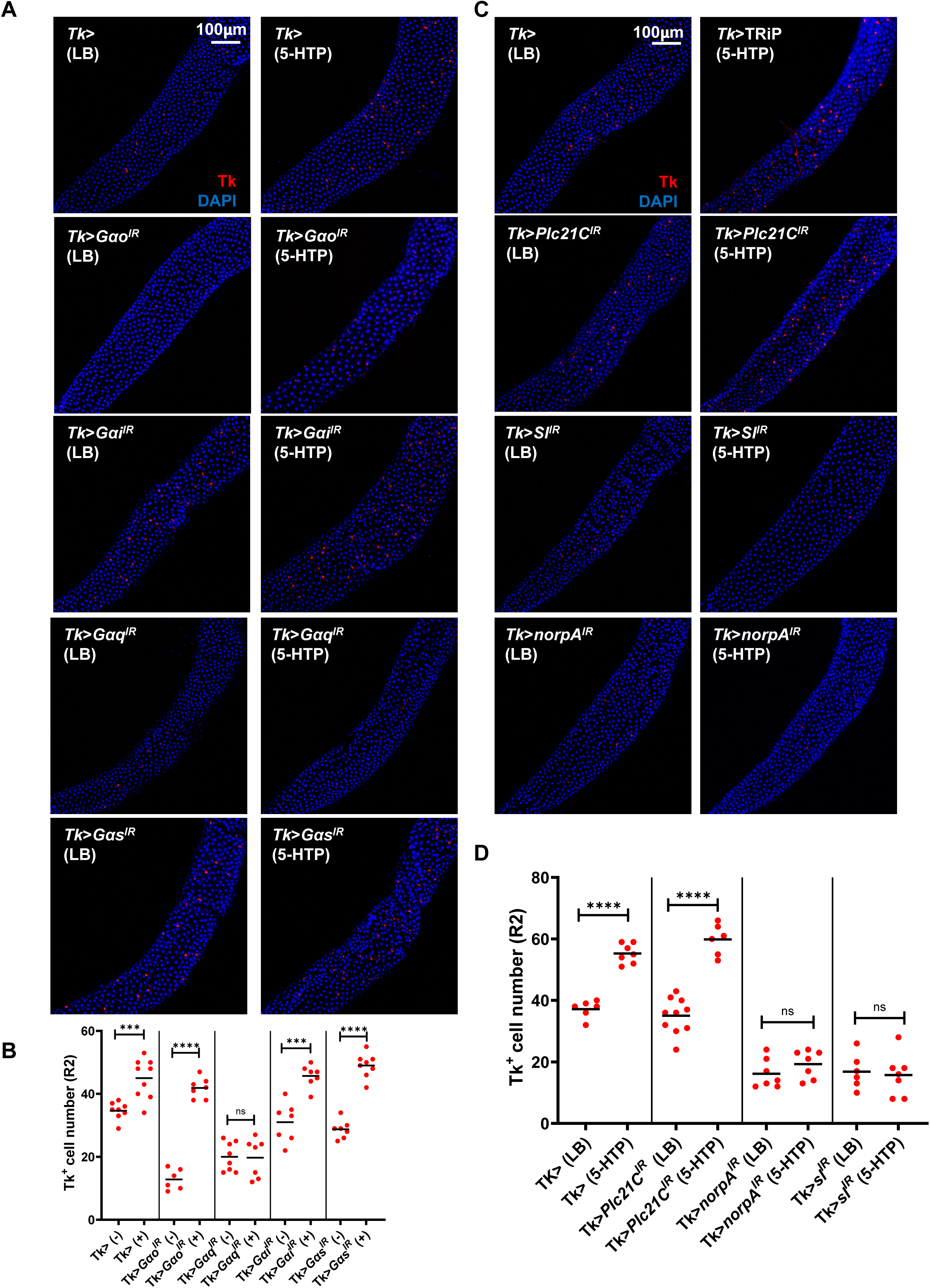
Serotonin receptors signal via Gα and phospholipase C paralogs no receptor potential (norpA) and small wing (sl) in response to 5-HT. (A) Immunofluorescent images and (B) quantification of Tk+ cells in the AMG (R2) of Tk> driver-only flies and the indicated G proteins knockdown flies fed LB alone or supplemented with 5-HTP (C) Immunofluorescent images and (D) quantification of Tk+ cells in the AMG (R2) of Tk> driver-only flies and the indicated phospholipase C paralog knockdown flies fed LB alone or supplemented with 5-HTP. In both experiments, a minimum of 6 intestines were visualized. The bar represents the mean. A Welch’s t test was used to assess significance. **** p<0.0001, *** p<0.001, ns not significant.

PLC cleaves phosphatidylinositol lipid species in the cell membrane to generate diacyl glycerol (DAG) and inositol triphosphate. DAG derived from membrane lipids promotes PKC’s relocation to the plasma membrane where it is activated and competent to phosphorylate protein targets. We hypothesized that PKC was downstream of PLC in the serotonin response pathway. To test this, we first fed OreR control flies the PKC inhibitor chelerythrine chloride (CC). Dietary supplementation with CC decreased Tk+ EECs and increased lipid accumulation in the AMG but not the PMG (Fig 5A and B and Fig S5A and B). We then tested the role of the six PKC paralogs, PKC53E, PKC98E, aPKC, Pkn, PKCδ, and inactivation no afterpotential C (inaC) in Tk+ EECs. Only Tk>*PKC53E*^RNAi^ and Tk>*PKC98E*^RNAi^ blocked 5-HTP-induced Tk expression and promoted lipid accumulation (Fig 5C-D and S5C-E).

**Figure 5:**
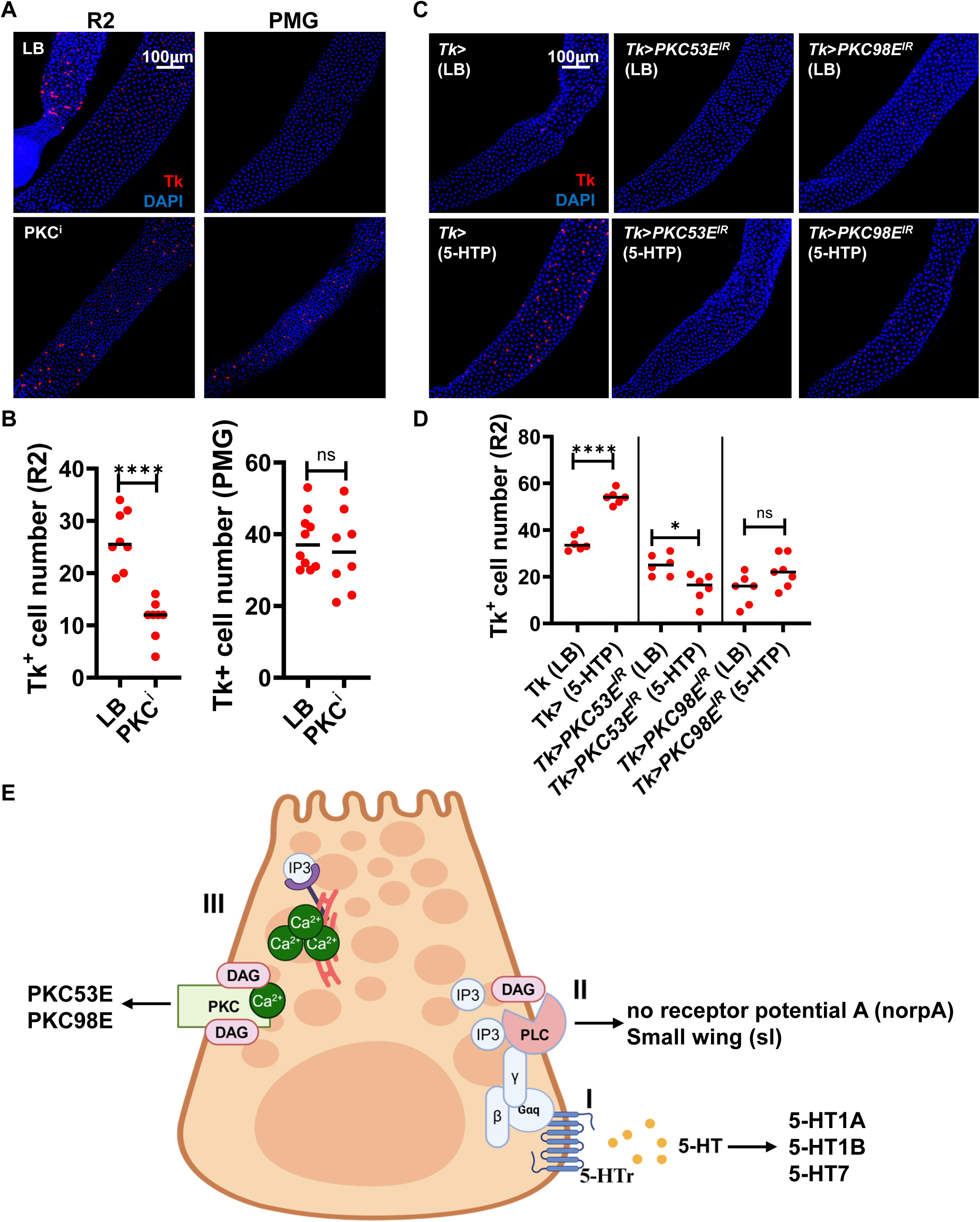
The serotonin signaling pathway in Tk+ cells activates protein kinase C paralogs PKC53E and PKC98E. (A) Immunofluorescent images and (B) quantification of Tk+ cells in the AMG (R2) and PMG of control flies fed either LB alone or supplemented with the protein kinase C inhibitor Chelerythrine Chloride (PKC^i^). (C) Immunofluorescent images and (D) quantification of Tk+ cells in the AMG (R2) of Tk> driver-only flies and the indicated protein kinase C knockdown flies fed LB alone or supplemented with 5-HTP. For (B) and (D), at least 6 intestines were examined. The bar represents the mean. A Welch’s t test was used to assess significance. **** p<0.0001, * P<0.01, ns not significant. (E) Schematic of the serotonin signaling pathway in Tk+ enteroendocrine cells.

The schematic shown in Fig 5E illustrates the pathway leading from serotonin receptor signaling to protein kinase C based on the results reported above. 5-HT receptors 1A,1B, and 7 respond to intestinal serotonin by interacting with Gαq which then activates the PLC homologs norpA and sl. These enzymes generate DAG in the cell membrane, which activates the protein kinase C homologs PKC53E and 98E. The next goal was to identify the target of these PKCs that regulates Tk and AMP expression.

### Serotonin promotes ecdysone-signaling in the gut

PKCs are serine/threonine kinases that promote the phosphorylation of substrates involved in diverse biological processes ^57^. The arthropod sex steroid 20-hydroxyecdysone (20E) has previously been shown to amplify IMD signaling by increasing transcription of the pattern recognition receptor PGRP-LC, and we showed that, in Tk+ EECs, this process regulates Tk transcription, lipid homeostasis, and AMP expression in the *Drosophila* intestine ^33, 34, 58, 59^. We first looked for evidence that the PKC target could be involved in ecdysone signaling by measuring transcription of shade (shd), which catalyzes the last step in 20E synthesis, and the 20E-induced proteins Eip74EF and Eip75B in the intestines of flies fed 5-HTP. As shown in Figure 6A, 5-HTP supplementation of OreR flies increased intestinal transcription of *Eip74EF* and *Eip75B* but not *shd* (Fig 6A). Furthermore, blocking serotonin synthesis in ECs with NP1>*Hn*^RNAi^ decreased transcription of *Eip74EF* and *Eip75B* (Fig 6B). As a final test, we used the ecdysone response element (EcRE)-driven *LacZ* (*EcRE-LacZ*) reporter to examine the impact of 5-HTP on overall 20E signaling activity in the gut. X-Gal staining showed that dietary supplementation with 5-HTP promoted intestinal ecdysone signaling (Fig 6C). As a positive control, 20E feeding also increased ecdysone signaling. We conclude that serotonin activates 20E signaling in the gut.

**Figure 6:**
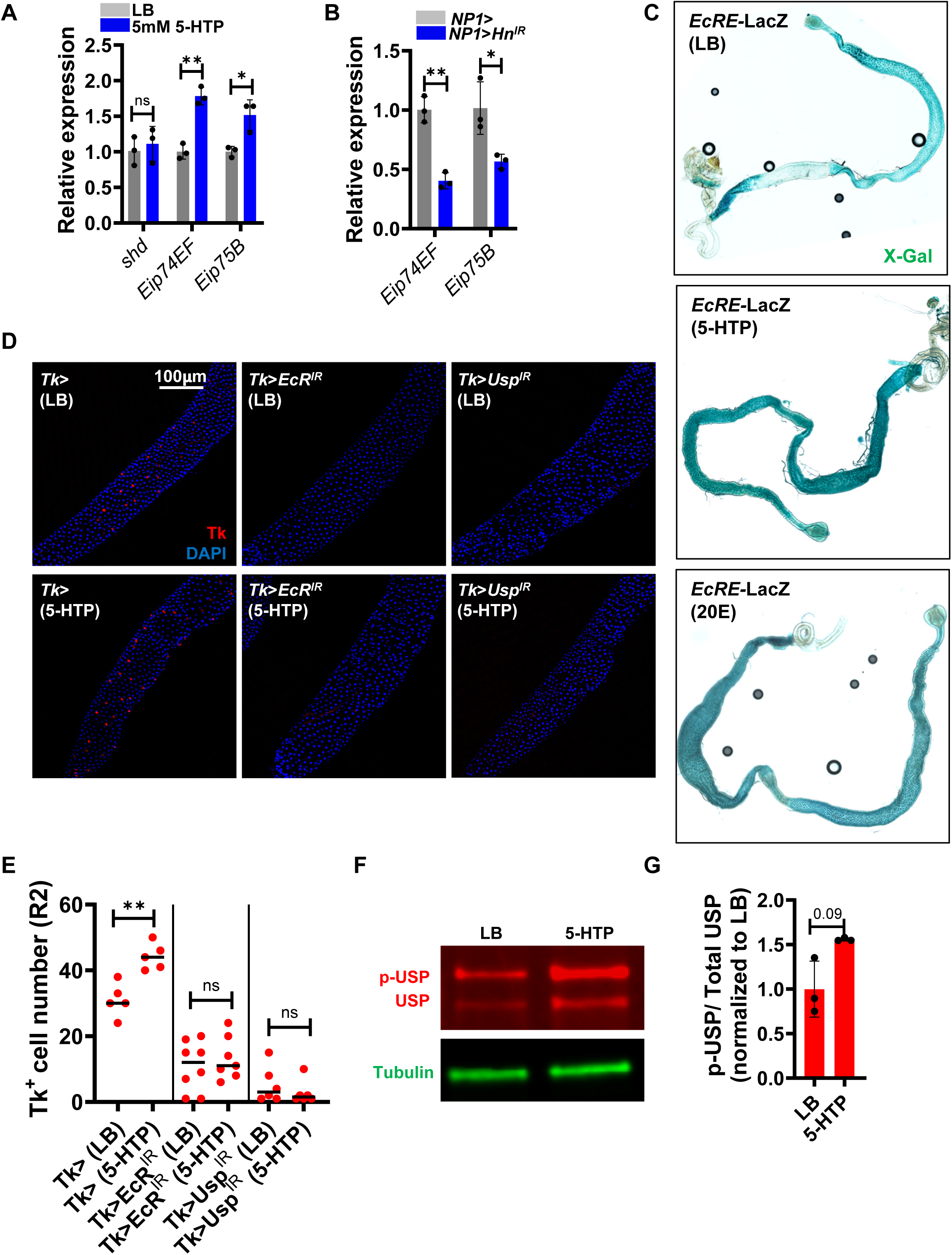
Serotonin amplifies transcription of ecdysone-regulated genes via phosphorylation of USP. RT-qPCR analysis of the indicated ecdysone-regulated genes (A) in the intestines of flies fed LB alone or supplemented with 5-HTP and (B) in control NP1> enterocyte driver-only and NP1>Hn^RNAi^ knockdown fly intestines. Bars show the mean of three biological replicates. Error bars represent the standard deviation. A student’s t test was used to assess significance. (C) The intestines of flies carrying an ecdysone response element (EcRE) fused to the gene encoding β-galactosidase that were fed either LB alone or supplemented with 5-HTP or 20-hydroxyecdysone (20E). Intestines were visualized after incubating with the β-galactosidase substrate 5-bromo-4-chloro-3-indolyl-β-D-galactopyranoside (X-Gal). (D) Immunofluorescence and (E) quantification of Tk+ cells in the intestines of Tk> driver-only flies and Tk>EcR^RNAi^ or Tk>Usp^RNAi^ flies fed LB alone or supplemented with 5-HTP. A minimum of 6 intestines were examined. Bars show the mean. A t test was used to assess significance. (F) Representative Western blot analysis and (G) quantification of phosphorylated USP in the intestines of flies fed LB alone or supplemented with 5-HTP. Tubulin was used as a loading control. Western blots were performed in triplicate. Bars show the mean. Error bars represent the standard deviation. A Welch’s t test was used to assess significance. ** p<0.01, * p<0.05, ns not significant.

### Activation of Tk and AMP expression by 5-hydroxytryptophan is dependent on ecdysone signaling and PGRP-LC

Our data show that 5-HTP activates ecdysone signaling, Tk expression, and AMP transcription in the intestine. We previously reported that 20E signaling activates the IMD pathway and Tk and AMP expression by increasing transcription of PGRP-LC in Tk+ EECs ^33^. Therefore, we first questioned whether the impact of serotonin on Tk expression was dependent on 20E signaling. To test this, we decreased expression of ultraspiracle (usp) and the ecdysone receptor (EcR) in Tk+ cells by RNAi ^60, 61^. Together, usp and EcR regulate 20E-dependent gene expression by binding to DNA as a heterodimer ^62^. In fact, Tk>*EcR*^RNAi^ and Tk>*usp*^RNAi^ blocked the increase in Tk+ EECs observed with dietary 5-HTP supplementation and resulted in lipid accumulation in the AMG (Fig 6D and E and Fig S6). This shows that 20E signaling is required for the serotonin-mediated increase in Tk+ EECs.

To examine whether either of the IMD-specific PRRs PGRP-LC or PGRP-LE were downstream of 5-HTP, we first measured PGRP-LC and PGRP-LE expression in the intestines of flies fed LB alone or supplemented with 5-HTP. As shown in Fig S7A, supplementation with 5-HTP increased expression of PGRP-LC but not PGRP-LE. We then questioned which of these PRRs was required for induction of AMP transcription in response to 5-HTP. Induction of AMPs was observed in a PGRP-LE*^112^* mutant but not the PGRP-LC^Δ5^ mutant (Fig S7B and C). In addition, PGRP-LC^Δ5^ abolished 5-HTP-induction of Tk+ EECs (Fig S7D and E). Taken together, we conclude that 5-HTP induces the IMD pathway by increasing intestinal transcription of *PGRP-LC* via 20E signaling.

### Dietary supplementation with 5-HTP increases phosphorylation of USP in the gut

There is evidence from the literature that usp phosphorylation by PKC homologs at Ser35 in S2 cells and salivary glands activates 20E signaling ^63^. We hypothesized that usp might be a phosphorylation target of PKCs in Tk+ cells as well. To test this, we fed flies 5-HTP and measured usp phosphorylation in the gut by Western blot analysis using an anti-usp antibody. As shown in Figure 6F-G, supplementation of fly diet with 5-HTP increased the ratio of phosphorylated-USP in the intestine. This suggests that serotonin promotes ecdysone signaling and Tk expression by increasing phosphorylation of usp.

### Serotonin and acetate are both required to activate IMD signaling in Tk+ cells

Here we present evidence that *V. cholerae*-produced tryptophan activates the IMD pathway by phosphorylating usp to promote 20E signaling. We previously reported that bacteria-derived acetate also increases 20E and IMD signaling via the action of the Tip60 histone acetyl transferase complex on variant histone 2A ^33,34^. We questioned whether just one of these microbial metabolites was sufficient to activate IMD signaling or whether both were required. Because the intestines of conventionally-raised flies are exposed to both tryptophan and acetate provided by commensal microbes, we used flies cured of commensal microbes by an antibiotic cocktail. These flies had decreased numbers of Tk+ EECs in their AMGs and accumulated lipids in their intestines, and both phenotypes could be rescued by dietary provision of 5-HTP (Fig 7A and B). In addition, 5-HTP increased transcription of ecdysone-regulated genes and IMD-regulated AMPs (Fig 7C and D). This suggests that in a control antibiotic-treated fly, supplementation of the diet with 5-HTP is sufficient to rescue the intestinal phenotype.

**Figure 7:**
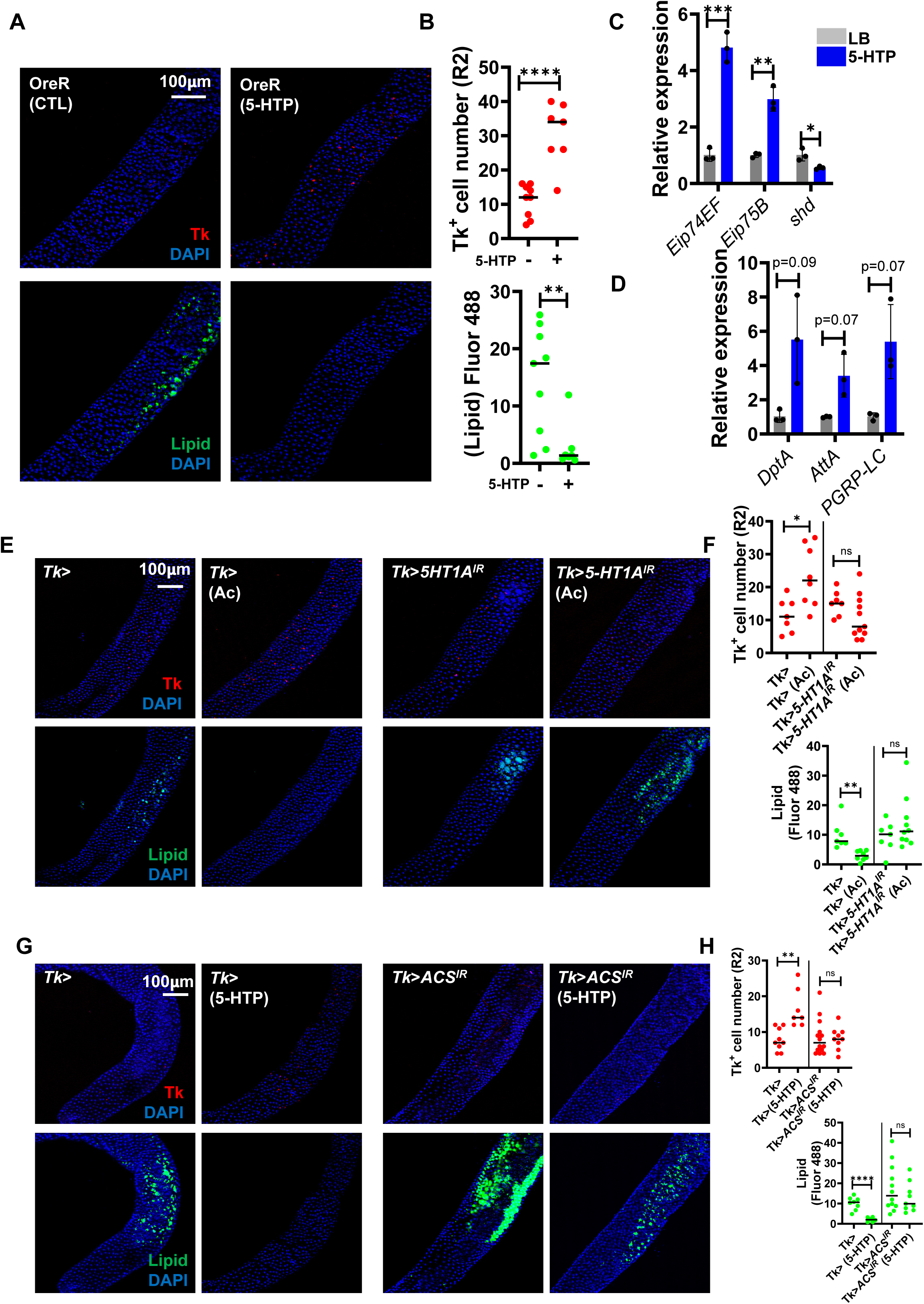
Both serotonin and acetate signals are required to for innate immune signaling via Tk+ cells. All experiments shown in this figure are performed with antibiotic-treated flies. (A) Fluorescent images and (B) quantification of Tk+ cells and lipid accumulation in the intestines of antibiotic-treated OreR flies fed either LB alone or supplemented with 5-HTP. A student’s t test was used to assess significance. RT-qPCR of (C) the indicated antimicrobial peptides, and (D) the indicated ecdysone-regulated genes in the intestines of antibiotic-treated flies fed LB alone or supplemented with 5-HTP. Bars show the mean of three biological replicates. Error bars represent the standard deviation. For (C), significance was assessed using a Welch’s t test. For (D), significance was assessed using a student’s t test. (E) Fluorescent images and (F) quantification of Tk+ cells and lipids in the intestines of antibiotic-treated Tk> driver-only flies or Tk>5-HT1A^RNAi^ flies fed LB alone or supplemented with acetate. Significant differences in Tk+ cells were evaluated using a student’s t test. For lipids, a Welcch’s t test was used. (G) Fluoresacent images and (H) quantification of Tk+ cells and lipids in the intestines of antibiotic-treated Tk> driver-only flies or Tk>ACS^RNAi^ flies fed LB alone or supplemented with 5-HTP. A student’s t test was used to assess significance. For all fluorescence experiments, a minimum of six intestines were examined. Bars represent the mean. **** p<0.0001, *** p<0.001, ** p<0.01, * p<0.05, ns not significant.

Acetate can be generated from pyruvate in mammalian cells and then converted into acetyl-CoA by acetyl-CoA synthase (ACS) ^64^. To determine whether acetate generated within EECs might be sufficient to restore intestinal homeostasis when 5-HTP is provided in the diet, we fed 5-HTP to Tk> driver and Tk>*ACS*^RNAi^ flies. Tk>*ACS*^RNAi^ completely blocked 5-HTP rescue (Fig 7G and H). We conclude that 5-HTP only rescues IMD signaling in Tk+ EECs when a basal level of acetate is present.

We then questioned whether serotonin signaling was essential to restore intestinal homeostasis when the diet of microbe-free flies was supplemented with acetate. To test this, we measured Tk+ EECs and intestinal lipid accumulation in the intestines of Tk> driver and Tk>*5-HT1A*^RNAi^, Tk>*5-HT1B*^RNAi^, and Tk>*5-HT7*^RNAi^ acetate-supplemented, antibiotic-treated flies. In each case, acetate was not sufficient to rescue Tk+ EECs and lipid homeostasis in the absence of the serotonin receptor (Fig 7E and F and Fig S8). We conclude that acetate and serotonin act synergistically to activate the innate immune response in Tk+ EECs and, furthermore, provision of acetate and tryptophan by intestinal microbes and/or the diet bolsters metabolic homeostasis in the arthropod anterior midgut.

## Discussion

Both commensal and pathogenic bacteria have evolved strategies to exploit rather than quell the host innate immune response. Here we show that *V. cholerae*, which colonizes the intestinal surface of environmental arthropods and mammals, secretes tryptophan into the intestine to activate the innate immune response by phosphorylating a nuclear steroid receptor. While this increases production of AMPs, it also augments synthesis of the host colonization factor Peri-15A, which is widely conserved in arthropods ^45^. We propose that this response has evolved to stabilize *V. cholerae’s* association with arthropod hosts.

Tryptophan is an essential amino acid that is provided in the laboratory fly diet in amounts that support rapid development. When a tryptophan-depleted diet is provided, the commensal microbiota can rescue the fly’s development by synthesizing and supplying tryptophan to the host ^49^. Furthermore, when *Hn* synthesis is blocked in uninfected flies, we observed decreased Tk+ EECs and lipid accumulation in enterocytes, suggesting that the diet and/or the microbiota provide sufficient tryptophan to maintain intestinal metabolic homeostasis. Because *V. cholerae* tryptophan secretion can further activate the host innate immune response, we conclude that this pathway is not saturated in uninfected flies.

Many animals detect the microbe-derived tryptophan metabolite indole via the aryl hydrocarbon receptor (AhR), which is expressed by intestinal epithelial and immune cells. AhR activation promotes the mucosal immune response through production of the pro-inflammatory cytokine interleukin (IL)-22 ^65, 66^. Microbes such as *Enterococcus gallinarum* and *Rodentibacter heylii*, which are found in the small intestine of neonatal mice, themselves produce serotonin from tryptophan. Serotonin synthesized by intestinal bacteria regulates host T cell development and promotes immune tolerance of commensal bacteria ^67^. Here we describe a novel role for microbe-derived tryptophan itself in manipulating the innate immune response of the model arthropod *Drosophila melanogaster*. This likely benefits the bacterium in that (i) it does not have to invest its own energy in serotonin synthesis and (ii) it co-opts a native host intercellular communication pathway that may deliver serotonin more efficiently to serotonin receptors on specific cell types.

Microbial metabolism is complex, and hosts must integrate responses to the many metabolites secreted by the intestinal microbiota ^68^. We previously showed that microbiota-derived acetate acts at the level of chromatin remodeling to increase ecdysone regulation, PGRP-LC transcription, and IMD signaling ^33, 34^. Here we describe a second microbial metabolite-responsive pathway that promotes ecdysone regulation through a distinct signaling cascade. Our results suggest that under homeostatic conditions, both metabolites are required to maintain metabolic homeostasis. At present, we have not established whether these pathways act synergistically to provide the amount of 20E signaling needed to maintain intestinal metabolic homeostasis or whether each is truly essential so that overactivating one pathway does not compensate for the absence of the other.

While researchers have focused on AMPs produced by PG-activated IMD signaling in ECs, the EEC IMD pathway responds uniquely to metabolites produced by bacteria such as acetate or tryptophan and produces EEC-specific effectors such as Tk and Peri-15a. We have shown that EEC-specific activation of Tk maintains intestinal lipid homeostasis, and we hypothesize that increased Peri-15A expression is one component in a branch of the innate immune response that reinforces the peritrophic matrix. Because they constitutively shed PG, microbes necessarily activate EC IMD signaling. However, they may also shape the intestinal innate immune response through secretion of metabolites that activate EEC IMD signaling. Here, we show that *V. cholerae* targets EEC-specific IMD signaling through tryptophan secretion to increase EEC expression of its colonization factor Peri-15a. We propose that this may represent a colonization strategy that stabilizes the interaction of this microbe with the arthropod intestinal epithelium.

Usp regulates 20E signaling by forming a heterodimer with its paralog, EcR. These two nuclear receptors are homologous to the androgen and estrogen nuclear receptors of mammals. Like these mammalian hormone receptors, EcR and usp are essential for developmental transitions but also play a role in adult animals ^58, 69, 70, 71^. Usp shares more identity and similarity with the estrogen receptors than it does with EcR and is most similar to the estrogen receptor β (Erβ), with 25% identity over 90% of its sequence. The phosphorylated serine of usp at position 35 is also conserved in Erβ. Erβ is expressed in the mammalian intestinal epithelium and is thought to impact inflammation ^72^. Our findings, therefore, provide a novel paradigm for microbial manipulation of intestinal inflammation that could extend to mammals.

*V. cholerae* is found in close contact with both terrestrial and aquatic arthropods in the environment. While these arthropods likely have differing microbiota that depend on their habitat and diet, the metabolism of most microbes including mixed acid fermentation and tryptophan synthesis is highly conserved. Furthermore, ecdysone and IMD signaling are highly conserved in both terrestrial and aquatic arthropod hosts. Therefore, it is plausible that the signaling pathway uncovered here that promotes *V. cholerae* colonization of the host intestine is an adaptation to maximize colonization of an environmental arthropod host.

## Acknowledgements

This work was supported by NIH R01AI158247 and NIH R01AI162701 to P.I.W. Anti-TK antibodies were generously provided by Jan Veenstra and AMP-deficient flies and the corresponding parental strain were generously provided by Bruno Lemaitre. The TK-Gal4 and NP1-Gal4 (Myo1A-Gal4) driver flies were kind gifts from Norbert Perrimon. Stocks obtained from the Bloomington Drosophila Stock Center (NIH P40OD018537) were used in this study. Microscopy images were acquired at the Microscopy Resources on the North Quad (MicRoN) core at Harvard Medical School. We thank Paola Montero Lopis at the MicRoN core for providing expertise with image acquisition and quantification.

## Experimental Methods

### Bacterial strains and growth conditions

Wild-type and Δ*trpE* mutant *V. cholerae* all derive from wild-type *V. cholerae* strain HC-494 isolated from a cholera patient in year 2013 of the Haitian *V. cholerae* epidemic. For all experiments, *V. cholerae* strains were culturing at 27°C in LB broth supplemented with 100 mg/mL streptomycin for 12-16 hours.

### Drosophila husbandry

Flies were raised on standard fly food containing 16.5 g/L yeast, 9.5 g/L soy flour, 71 g/L cornmeal, 5.5 g/L agar, 5.5 g/L malt, 7.5% corn syrup, and 0.4% propionic acid at 25 °C in a humidified incubator programmed to keep a 12 hr light/dark cycle. 5-7 days old female flies were used for all experiments. The following stocks were obtained from the Bloomington Drosophila Stock Center (BDSC): Ore R-C (BL 5), TRiP (BL36303), *Hn^IR^* (BL29540), *5-HT1A^RNAi^* (BL25834), *5-HT1B^RNAi^* (BL25833), *5-HT2A^RNAi^* (BL31882), *5-HT2B^RNAi^* (BL25874), *5-HT7^RNAi^* (BL27273), *Gαo^RNAi^* (BL28010), *Gαi^RNAi^*(BL40890), *Gαq^RNAi^* (BL33765), *Gαs^RNAi^* (BL29576), *Plc21C^RNAi^* (BL31269), *Sl^RNAi^* (BL32385), *norpA^RNAi^*(BL31113), *PKC53E^RNAi^* (BL27491), *PKC98E^RNAi^*(BL29311), *aPKC^RNAi^* (BL25946), *Pkn^RNAi^* (BL28335), *PKCδ^RNAi^* (BL28355), *inaC^RNAi^* (BL31651), *EcRE-LacZ* (BL4516), *EcR^RNAi^* (BL50712), *Usp^RNAi^* (BL27258), *PGRP-LC^Δ5^* (BL36323), *PGRP-LE^112^* (BL33055), *ACS^RNAi^*

(BL41917). The *NP1-Gal4 (Myo1A-Gal4)* and *Tk-Gal4* driver flies were kind gifts from Norbert Perrimon. 10 AMPs deleted (ΔAMP) and parental(*^W1118^*) fly was kind gift from Burno Lemaitre.

### Tryptophan (Trp), 5-hydroxytryptophan (5-HTP), 20-hydroxyecdysone (20E), and Chelerythrine chloride Supplementation

For oral Trp, 5-HTP and 20E treatment, ten 5- to 7-day-old female flies were placed in vials containing a cellulose acetate plug infused with 3 mL of L.B broth alone or supplemented with 5mM L-Tryptophan (Sigma-Aldrich T0254-100G), 5mM L-5-Hydroxytryptophan (Acros Organics 148290050),10µM 20E (Sigma-Aldrich H5142-5MG), or 100µM C.C (Abcam AB120600) for 3 days. The fly midguts were then dissected for microscopic or RT-qPCR analysis.

### *V. cholerae* infection and colonization assays

Oral Vibrio cholerae colonization assays were carried out in an arthropod containment level 2 facility. 10-15 female flies per genotype (5-7 days old) were infected with the *Vibrio cholerae* HC494 strain collected from Haiti in 2013 ^73^. Flies were placed in fly vials containing a cellulose acetate plug infused with a 3mL suspension of an overnight *Vibrio cholerae* culture diluted 1:10 in LB broth. For RT-qPCR and immunofluorescence experiments, midguts were harvested after exposure to *V. cholerae* for 3 days. For colonization assays, flies were fed with *V. cholerae* as above for two days and then placed on sterile PBS for one day.

### Immunofluorescence

The midguts of 7- to 10-day-old female flies were dissected and fixed in PBS containing 4% formaldehyde at room temperature for 20 minutes. After fixation, tissues were washed with PBS-T (PBS + 0.1% Tween 20) three times for 10 minutes at room temperature. After removing PBS-T, tissues were incubated in blocking solution (PBS-T with 2% BSA and 0.1% Triton X-100) for 1 hour at room temperature. Subsequent incubation of the gut with primary antibodies in blocking solution was performed overnight at 4°C. Primary antibodies used in this study were Rabbit anti-Tk (1:500, kind gift from Jan Adrianus Veenstra), Mouse anti-Prospero (MR1A)-c (1:100, Developmental Studies Hybridoma Bank AB_528440), Rat anti-Serotonin (1:300 Abcam AB6336). Samples were then washed three times for 10 minutes in PBS-T and incubated in blocking solution containing secondary antibody, DAPI (5µg/ml here, Invitrogen D1306) and BODIPY 493/503 (1µg/ml, Invitrogen D3922) for 2 hours at room temperature. Secondary antibodies used in this study were Alexa 594 conjugated anti-rabbit secondary antibody (1:500, Thermo Fisher Scientific A-11012), Alexa 488 conjugated anti-mouse secondary antibody (1:500, Thermo Fisher Scientific A-11001), and Alexa 488 conjugated anti-rat secondary antibody (1:500, Thermo Fisher Scientific A-11006). Samples were then washed again three times for 10 minutes in PBS-T. The tissues were mounted in mounting medium (Vector Laboratories H-1000). Confocal images were acquired using a Zeiss LSM 980 confocal microscope and 10X objective. Tachykinin-positive cells in R2 region of anterior midgut were counted manually. ImageJ was used to quantify fluorescence. This was normalized by dividing by the total area quantified. Background fluorescence was subtracted from this measurement. A minimum of 6 midguts were quantified for each condition and genotype.

### RT-qPCR analysis

Total RNA was extracted from a minimum of 10 midguts using TRIzol reagent (Thermo Fisher Scientific 15596026), followed by further purification using the Direct-zol RNA MiniPrep Plus kit (Zymo Research R2070), according to the supplier’s instructions. RNA was quantified using a NanoDrop 1000 Spectrophotometer. cDNA was synthesized from 500 ng of total RNA using the iScript cDNA Synthesis Kit (Bio-Rad 1708891) as described by the manufacturer. RT-qPCR was performed using iTaq SYBR Green (Bio-Rad 1725121) on a QuantStudio™ 3 or QuantStudio™ 5 real-time PCR system (Applied Biosystems). Three biological replicates were performed for each experiment, and duplicate technical measurements were included for each of the two experimental replicates. Relative expression of target genes was calculated using the comparative CT method normalized to *rp49* for *Drosophila* and *clpX* for *V.cholerae*. The primers used for RT-qPCR analysis are listed in Table S1.

### X-Gal staining

For X-Gal staining, the Senescence β-Galactosidase Staining Kit (Cell Signaling Technology #9860) was used according to supplier protocol as follows. Fly guts fixed as described above were washed three times with PBS for 10 minutes at room temperature. After removing PBS, tissues were incubated in 300µl β-Galactosidase Staining Solution at 37°C overnight or for 24 hours. After incubation, the tissues were mounted in 70% glycerol. Intestines were imaged using Olympus VS200 Slide scanner using 10X objective.

### Generation of Microbe-depleted Animals

For gut microbiota depletion, female adult flies aged 3- to 5-days-old were distributed into fly food containing an antibiotic cocktail, consisting of 500 mg mL^-1^ Ampicillin (Thermo Fisher Scientific 69-52-3), 50 mg mL^-1^ Tetracycline (IBI Scientific 64-75-5), 200 mg mL^-1^ Rifampicin (Calbiochem 557303), and 100 mg mL^-1^ Streptomycin (Sigma-Aldrich 3810-74-0) for 5 days, and then transferred to fly food containing an antibiotic cocktail, with or without 5mM 5-HTP, 50mM sodium acetate (Thermo Fisher Scientific 127-09-3) for 3 days. Flies were flipped to a fresh fly food every day.

### Quantification of commensal *Acetobacter* and *Lactobacillus sp*. in the *Drosophila* intestine

8 Female flies per condition were washed with 70% ethanol and PBS to lyse bacteria attached to the external surface of the fly and then homogenized. Serial dilutions of the homogenates were plated on deMan, Rogosaand Sharpe (MRS) agar plates (Sigma-Aldrich 69966). Colonies of Acetobacter and Lactobacillus sp. were identified by morphology, and the number of colony forming units (CFU) per fly was calculated.

### Western Blot Analysis

The intestines of ten female flies were dissected and homogenized with a pestle in 100µl PBS. After adding 25µl 4X Laemmli buffer with β-Mercaptoethanol, the homogenized fly intestines were incubated at 95°C for 10min. Protein was then separated on 4-20% polyacrylamide gel (BioRad 456-1096) and transferred to a PVDF membrane (BioRad 1704272). The membrane was blocked in Odyssey Blocking Buffer (LI-COR 927-60001) for 1 hour at 4°C with gentle shaking. Incubation of the membrane with primary antibodies in Blocking Buffer was performed overnight at 4°C. Primary antibodies used in this study were Rabbit anti-Ultraspiracle (1:1000, Abcam AB106341), mouse anti-β-tubulin (1:1000, Developmental Studies Hybridoma Bank AB_579794. Membranes were washed in TBS-T (TBS + 0.1% Tween 20) three times for 10 minutes and incubated in the Blocking Buffer containing secondary antibody for 2 hours at room temperature and then washed again in TBS-T three times for 10 minutes. Images were acquired using the Azure c600 imager. Secondary antibodies used in this study were IRDye 680RD Goat anti-Rabbit IgG Secondary Antibody (1:10000, LI-COR 926-68071), IRDye 800CW Goat anti-Mouse IgG Secondary Antibody (1:10000, LI-COR 926-32210).

### Statistical analysis

GraphPad Prism 10.0.3 software was used for statistical analyses and graph generation. Measurements represent the mean of at least three biological replicates in all graphs. Error bars represent mean ± the standard deviation. As appropriate, a two-tailed Student’s t test, ordinary one-way ANOVA or Brown-Forsythe ANOVA with post hoc Dunnett’s test was used to calculate significance. The statistical test applied is noted in the Figure Legend. For all tests, statistical significance was calculated as a p value. Asterisks indicate the following p value ranges: * for p < 0.05, ** for p < 0.01, *** for p < 0.001, **** for p < 0.0001, and ns (non-significant) for p > 0.05.

## Acknowledgements

This work was supported by NIH R01AI158247 and NIH R01AI162701 to P.I.W. Anti-TK antibodies were generously provided by Jan Veenstra, and AMP-deficient flies and the corresponding parental strain were generously provided by Bruno Lemaitre. The TK-Gal4 and NP1-Gal4 (Myo1A-Gal4) driver flies were kind gifts from Norbert Perrimon. Stocks obtained from the Bloomington Drosophila Stock Center (NIH P40OD018537) were used in this study. Microscopy images were acquired at the Microscopy Resources on the North Quad (MicRoN) core at Harvard Medical School. We thank Paola Montero Lopis at the MicRoN core for providing expertise with image acquisition and quantification.

## Author Contributions

X. D. and P. I. W. designed the experiments. X. D. performed the experiments. X. D. and P. I. W. analyzed the data. X. D. wrote the manuscript. All authors reviewed, edited, and approved the manuscript.

## Supplementary Figure Legends

**Figure S1:**
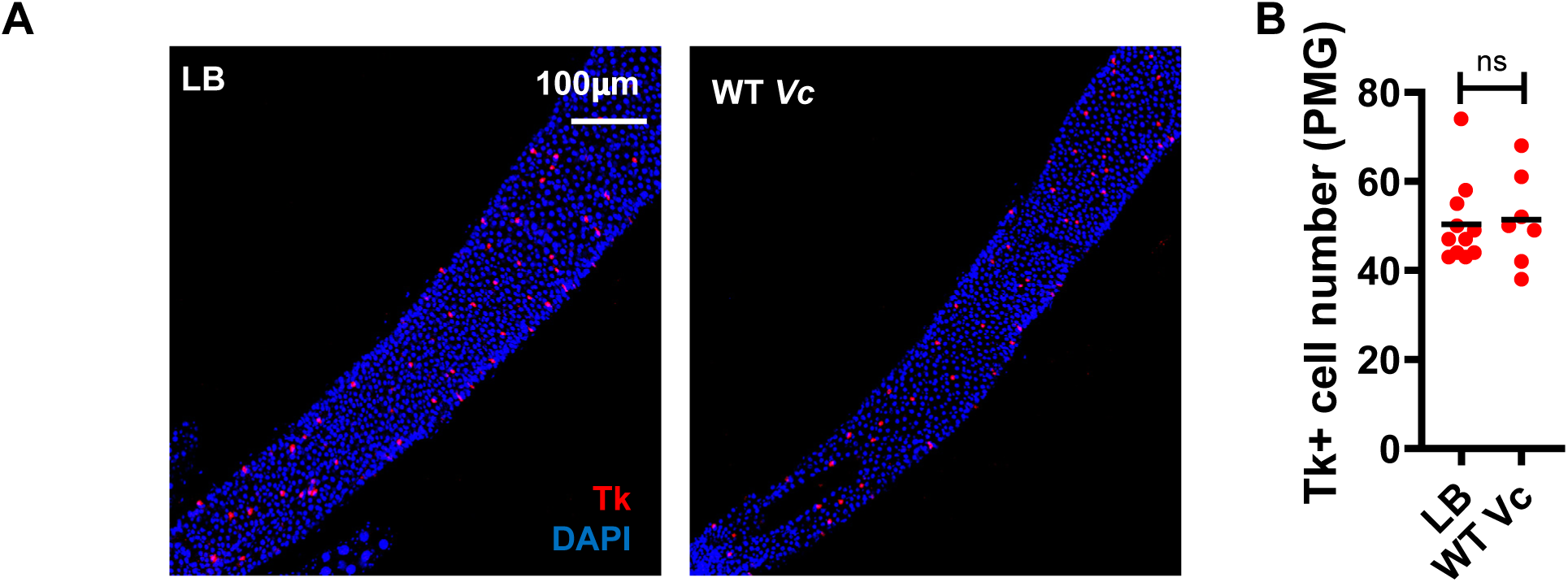
*V. cholerae* infection does not alter Tachykinin (Tk)+ cell numbers in the posterior midgut. Related to Fig 1. (A) Immunofluorescence and (B) quantification of Tk+ cells in the posterior midgut of OreR flies fed LB alone or inoculated with wild-type *Vibrio cholerae* (WT Vc). The mean of at least six intestines is shown. A student’s t test was used to assess significance. ns not significant.

**Figure S2:**
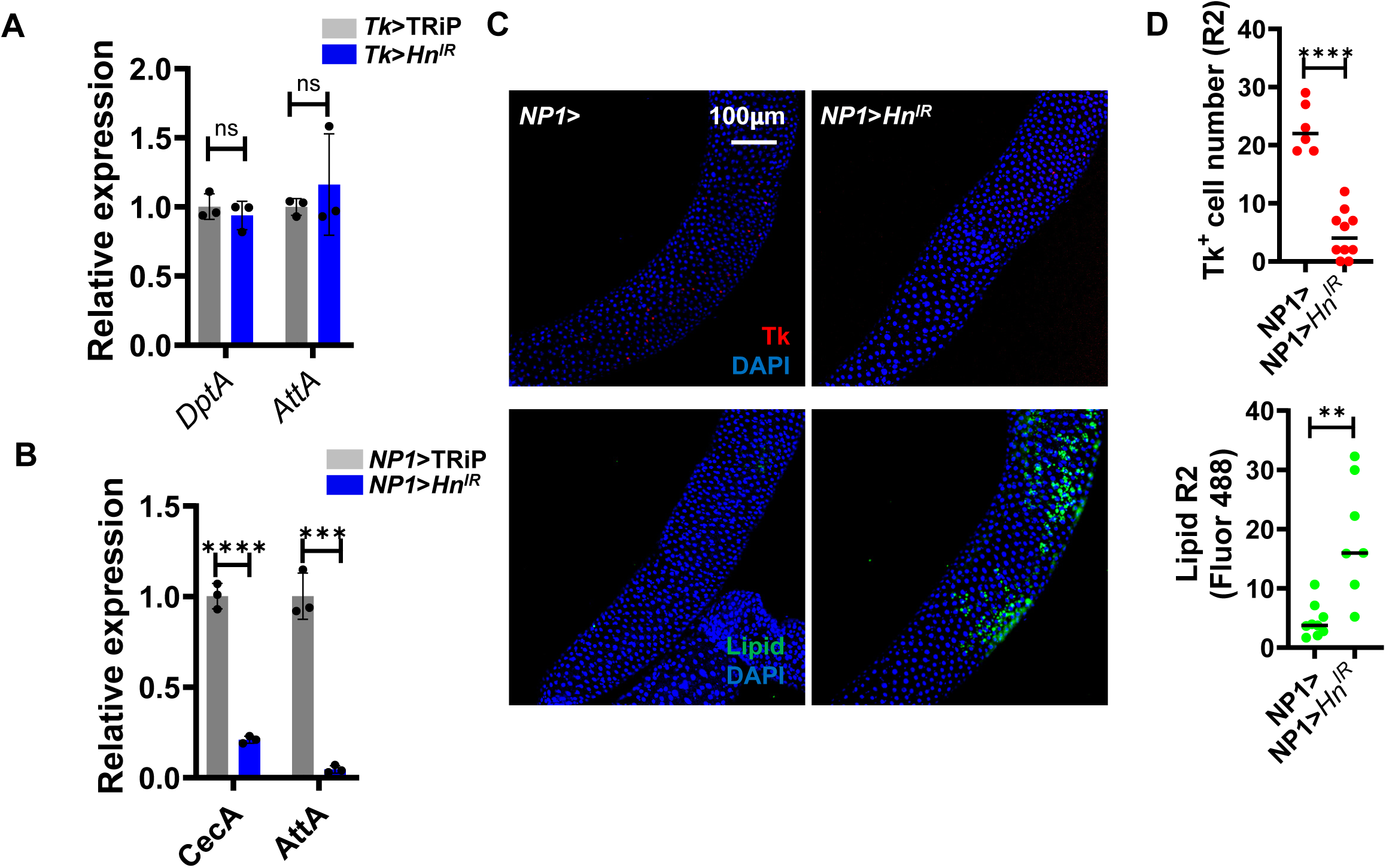
Serotonin synthesis in ECs of uninfected flies impacts IMD signaling and Tk expression. (A) RT-qPCR analysis of the antimicrobial peptide genes *CecA1* and *AttA* in the intestines of enterocyte NP1> driver-only flies and NP1>Hn^RNAi^ flies with decreased transcription of tryptophan hydroxylase. (B) RT-qPCR analysis of the antimicrobial peptide genes *DptA* and *AttA* in the intestines of Tk-expressing enteroendocrine cell Tk> driver-only flies and Tk>Hn^RNAi^ flies. The mean of biological triplicates is shown. Error bars represent the standard deviation. (C) Fluorescent images and (D) quantification of Tk+ cells and lipid accumulation in the R2 region of the intestines of flies described in (A). The mean of at least six intestines is shown for each condition. For quantification of Tk+ cells, a student’s t test was used to assess significance. A Welch’s t test was used for lipid quantification analysis. **** p<0.0001, *** p<0.001, ** p<0.01, ns not significant.

**Figure S3:**
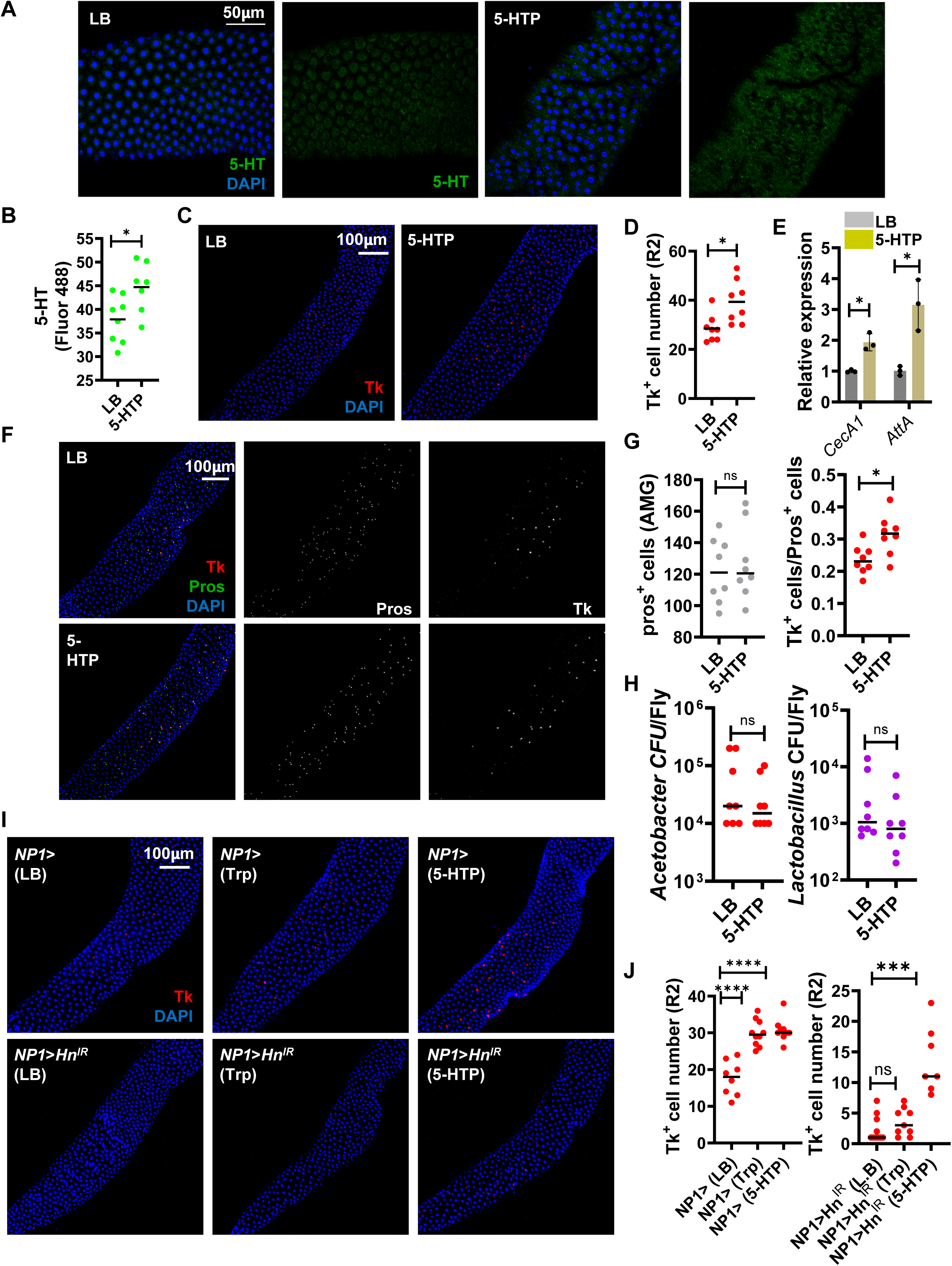
The phenotype of 5-HTP-supplemented flies is similar to that of *V. cholerae* infected flies. Immunofluorescent images and quantification of (A-B) serotonin and (C-D) Tk+ cells in the R2 region of the AMG Oregon R control flies fed either LB alone or supplemented with 5 mM 5-hydroxytryptophan (5-HTP). The brightness of the 5-HT micrographs was similarly corrected to improve visualization. The mean of at least six intestines is shown for each condition. A student’s t test was used to assess significance. (E) RT-qPCR analysis of *CecA1* and *AttA* expression in Oregon R control flies fed either LB alone or supplemented with 5-HTP. Bars represent the mean of three biological replicates. A Welch’s t test was used to assess significance. (F) Immunofluorescent images and (G) quantification of Prospero (Pros)+ cells and Tk+ cells in the R2 region of the intestines of flies fed LB alone or supplemented with 5-HTP. The mean of at least six intestines is shown for each condition. A student’s t test was used to assess significance. (H) Measurement of colony forming units (CFU) of *Acetobacter* and *Lactobacilli* in the intestines of Oregon R control flies fed either LB alone or supplemented with 5-HTP. The mean of 8 flies is shown. A Welch’s t test was used to assess significance. (I) Immunofluorescent images and (J) quantification of Tk+ cells in the AMG of enterocyte driver-only NP1> or NP1>Hn^RNAi^ flies fed LB alone or supplemented with tryptophan (Trp) or 5-HTP. The mean of the intestines of at least six flies is shown. A ordinary one-way ANOVA with Dunnett’s multiple comparisons test was used to assess significance **** p<0.0001, *** p<0.001, * p<0.05, ns not significant.

**Figure S4:**
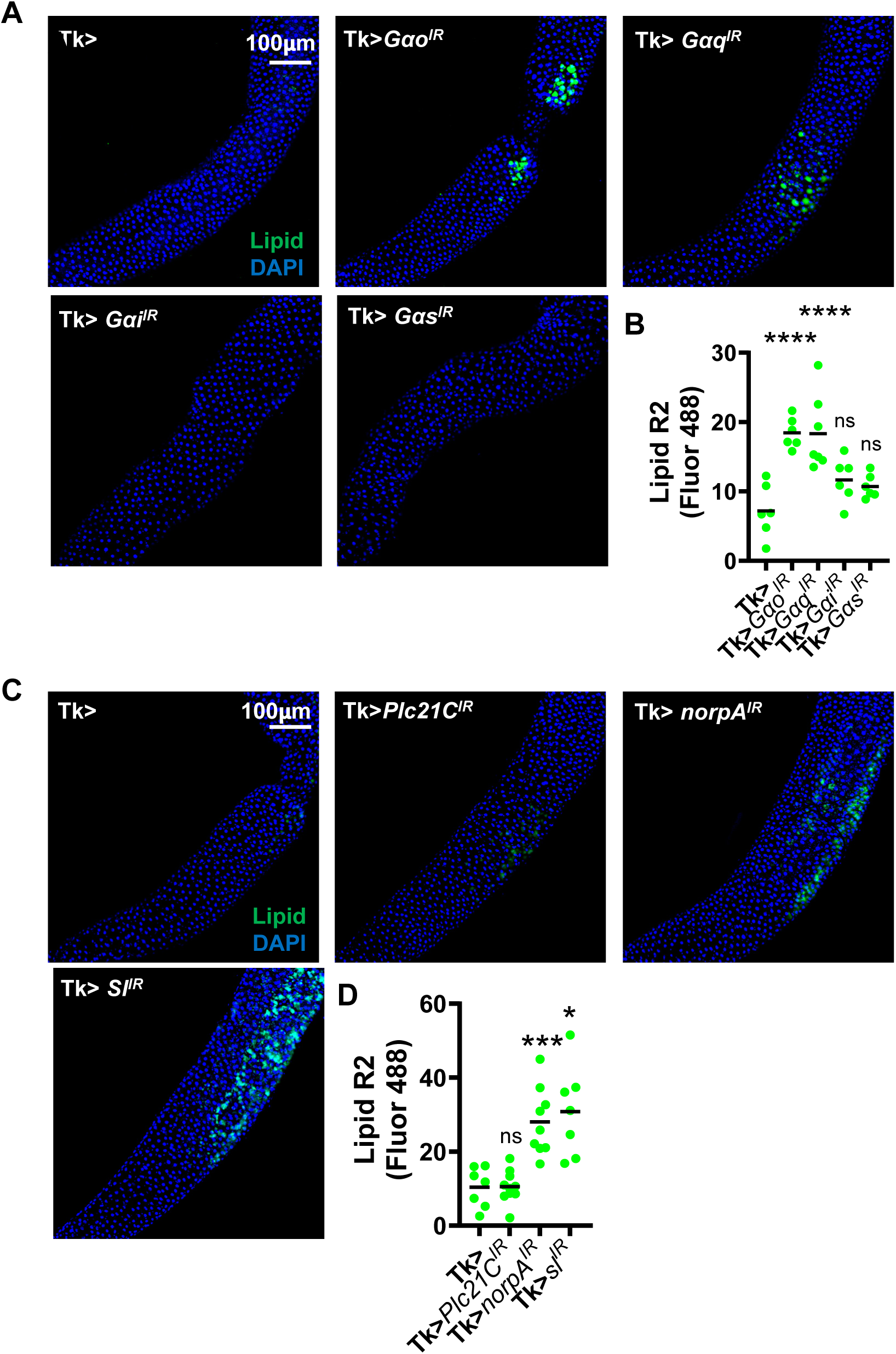
Knockdown of Gαq and phospholipase C paralogs no receptor potential (norpA) and small wing (sl) leads to intestinal lipid accumulation. (A) Lipid staining with Bodipy and (B) quantification of fluorescence in the AMG of Tk> driver-only flies and the indicated G protein knockdown flies. (C) Immunofluorescent images and (C) Lipid staining with Bodipy and (D) quantification of fluorescence in the AMG of Tk> driver-only flies and the indicated phospholipase C paralog knockdown flies. In both experiments, a minimum of 6 intestines were visualized. The bar represents the mean. In (B), an ordinary one-way ANOVA with a Dunnett’s multiple comparisons test was used to assess significance. In (D), a Welch’s and Brown-Forsythe ANOVA with a Dunnett’s T3 multiple comparisons test was used. **** p<0.0001, * p<0.05, ns not significant.

**Figure S5:**
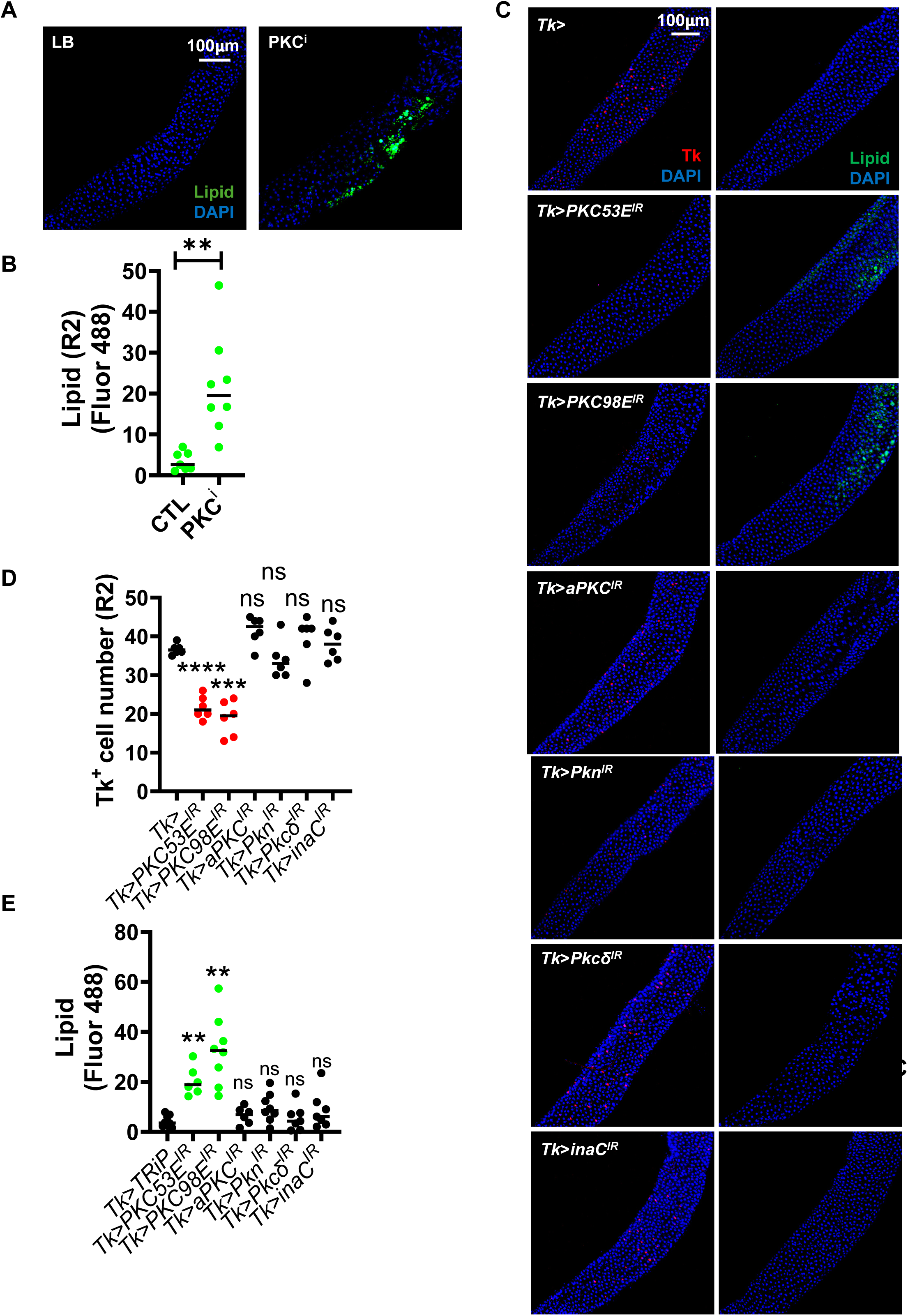
A protein kinase C inhibitor and knockdown of PKC53E and PKC98E leads to intestinal lipid accumulation. (A) Lipid staining with Bodipy and Tk immunofluorescence and quantification (B) in the AMG of OreR flies fed LB alone or supplemented with the protein kinase C inhibitor chelerythrine chloride (PKC^i^). A Welch’s t test was used to assess significance. (C) Immunofluorescence staining of Tk+ cells and Bodipy lipid staining in the AMG of Tk> driver-only flies and the indicated protein kinase C knockdown flies. (D) Quantification of Tk+ cells and (E) quantification of Bodipy fluorescence in the AMG of the flies described in (C). For (D-E), the mean of at least 6 intestines is shown. A Welch’s and Brown-Forsythe ANOVA with a Dunnett’s T3 multiple comparisons test was used to assess significance. **** p<0.0001, *** p<0.001, ** p<0.01, ns not significant.

**Figure S6:**
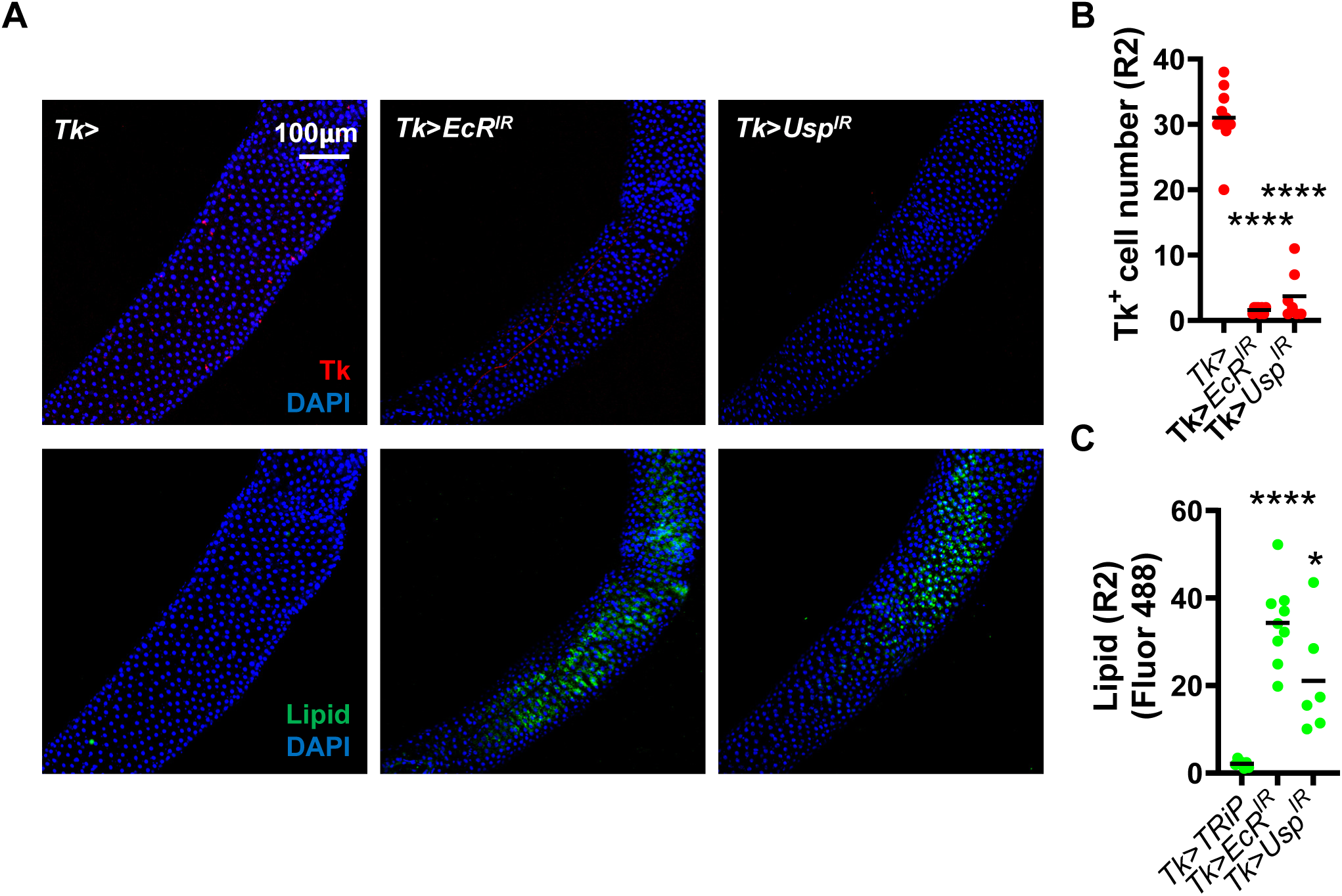
Knockdown of EcR and USP in Tk+ cells results in decreased Tk immunofluorescence and intestinal lipid accumulation. (A) Lipid staining with Bodipy and Tk immunofluorescence and quantification (B) of Tk+ cells and (C) Bodipy fluorescence in the AMG of Tk-expressing enteroendocrine cell Tk> driver-only or Tk>EcR^RNAi^ and Tk>Usp^RNAi^ knockdown flies. The mean of at least 6 intestines is shown. A Brown-Forsythe and Welch’s ANOVA with a Dunnett’s T3 multiple comparisons test was used to assess significance. **** p<0.0001, * p<0.01, ns not significant.

**Figure S7:**
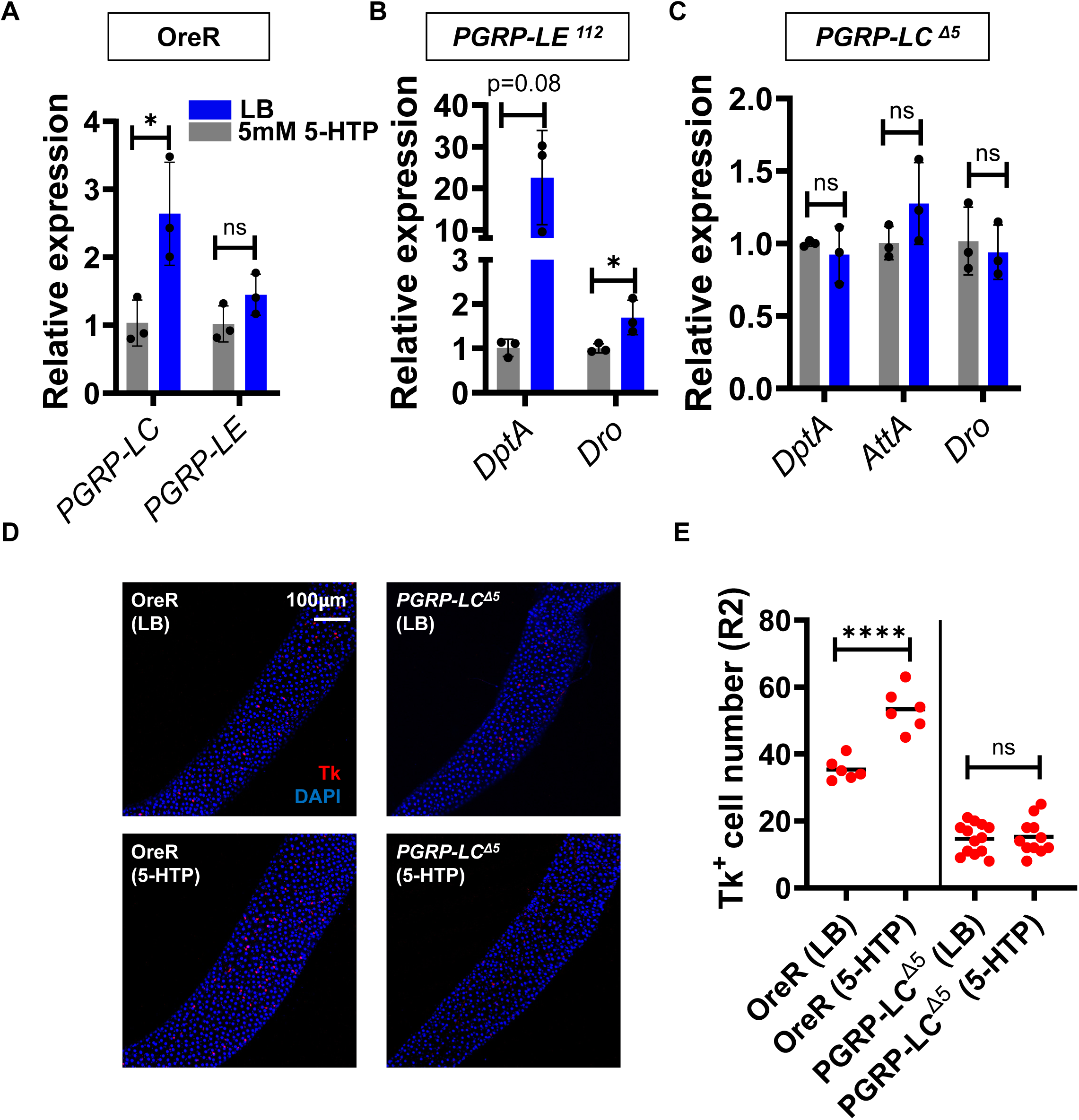
Evidence that 5-HTP activates the intestinal innate immune response by increasing transcription of PGRP-LC. RT-qPCR of (A) *PGRP-LC* and *PGRP-LE* expression in control OreR and the indicated antimicrobial peptide genes in (B) PGRP-LE^112^ mutant, and (C) PGRP-LC^Δ5^ mutant flies fed LB alone or supplemented with 5-HTP. The mean of biological triplicates is shown. Error bars represent the standard deviation. A student’s t test was used to assess significance for all measurements except for *DptA* in (B), where a Welch’s t test was used. Immunofluorescence (D) and quantification (E) of Tk+ cells in the AMG (R2) of OreR control or PGRP-LC^Δ5^ flies fed LB alone or supplemented with 5-HTP. The mean of at least 6 intestines is shown. A student’s t test was used to assess significance. **** p<0.0001, * p<0.05, ns not significant.

**Figure S8:**
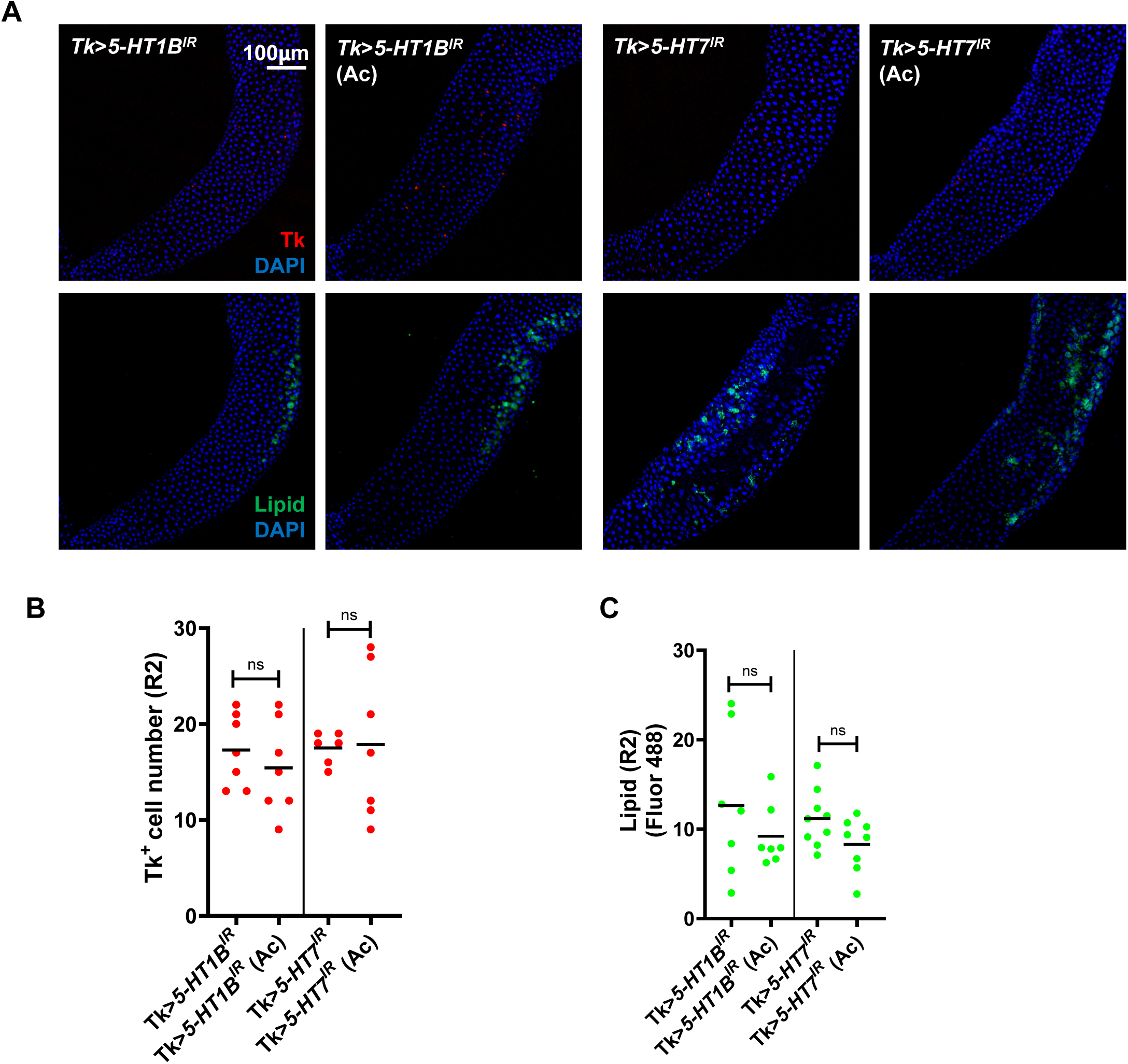
Serotonin receptor knockdown in Tk-expressing cells prevents acetate rescue of the germ-free phenotype. All experiments shown in this figure are performed with antibiotic-treated flies. (A) Tk immunofluorescence and lipid staining with Bodipy and quantification of (B) Tk+ cells and (C) Bodipy fluorescence in the AMG of enteroendocrine cell Tk> driver-only or Tk>*5-HT1A*^RNAi^ and Tk>*5-HT7*^RNAi^ flies with or without acetate (Ac) supplementation of the diet. A minimum of six intestines was examined. Bars represent the mean. A student’s t test was used to assess significance ns not significant.

**Table S1: Primers used for qRT-PCR analysis of gene transcription in *Drosophila melanogaster* and *Vibrio cholerae*.**

